# Bacterial carotenoids suppress *Caenorhabditis elegans* surveillance and defense of translational dysfunction

**DOI:** 10.1101/2020.01.08.898668

**Authors:** J. Amaranath Govindan, Elamparithi Jayamani, Victor Lelyveld, Jack Szostak, Gary Ruvkun

## Abstract

Microbial toxins and virulence factors often target the eukaryotic translation machinery. *Caenorhabditis elegans* surveils for such microbial attacks by monitoring translational competence, and if a deficit is detected, particular drug detoxification and bacterial defense genes are induced. The bacteria *Kocuria rhizophila* has evolved countermeasures to animal translational surveillance and defense pathways. Here, we used comprehensive genetic analysis of *Kocuria rhizophila* to identify the bacterial genetic pathways that inhibit *C. elegans* translational toxin surveillance and defense. *Kocuria rhizophila* mutations that disrupt its ability to disable animal immunity and defense map to multiple steps in the biosynthesis of a 50-carbon bacterial carotenoid from 5 carbon precursors. Extracts of the C_50_ carotenoid from wild type *K. rhizophila* could restore this bacterial anti-immunity activity to *K. rhizophila* carotenoid biosynthetic mutant. *Corynebacterium glutamicum,* also inhibits the *C. elegans* translation detoxification response by producing the C_50_ carotenoid decaprenoxanthin, and *C. glutamicum* carotenoid mutants are defective in this suppression of *C. elegans* detoxification. Consistent with the salience of these bacterial countermeasures to animal drug responses, bacterial carotenoids sensitize *C. elegans* to drugs that target translation and inhibit food aversion behaviors normally induced by protein translation toxins or mutations. The surveillance and response to toxins is mediated by signaling pathways conserved across animal phylogeny, suggesting that these bacterial carotenoids may also suppress such human immune and toxin responses.

## Introduction

Many bacterial toxins and virulence factors target the highly conserved RNAs and proteins of the ribosome and associated translation factors. In response to such toxin attacks, eukaryotic defense responses include the induction of specific suites of drug detoxification and anti-bacterial immunity genes. The textbook view is that these detoxification responses are triggered by chemical recognition or virulence factor receptor proteins (guard proteins) that then couple to signaling pathways for the induction of immunity and detoxification gene responses. Such a toxin recognition surveillance system can be defeated by the evolution of novel bacterial toxins with distinct chemical or protein sequence signatures. But a system that surveils for toxins by their toxicity to, for example, the eukaryotic system targeted, for example, translation of proteins, may be superior for the detection of novel bacterial toxins. The nematode *Caenorhabditis elegans* surveils a wide range of core cellular pathways for such deficits, and whether it is caused by a toxin, a mutation or RNA interference, detoxification and immunity genes are induced (Melo and Ruvkun, 2012)(Govindan et al., 2015). For example, *C. elegans* mutations in translation factor genes themselves trigger the same immune and detoxification gene inductions as those induced by bacterial toxins and virulence factors that inhibit *C. elegans* translation (Govindan et al., 2015). Thus, translation in *C. elegans* is monitored so that decrements in protein synthesis caused by toxins or mutations are detected and coupled to a defense response. The detected translational deficits are coupled via a conserved p38MAPK, bZIP/ZIP-2 transcription factor and the bile acid biosynthetic pathways to activate xenobiotic detoxification and bacterial immune response genes as well as food aversion behaviors (Melo and Ruvkun, 2012)(Govindan et al., 2015). By monitoring decrements in core cellular functions rather than the molecular structure of an unknown toxin, *C. elegans* can detect unanticipated pathogens and toxins. Many of the components of this signaling cascade, for example, the MAP kinase and bile acid biosynthetic pathway, are conserved across animals, suggesting that this system of toxin surveillance and response may not be parochial to *C. elegans*.

Such an animal defensive strategy may drive the evolution of bacterial pathogen countermeasures to thwart these defense responses (Melo and Ruvkun, 2012)(Samuel et al., 2016). Commensal bacteria may also seek to silence such animal defense responses to establish a benign or symbiotic relationship. Because bacteria synthesize a wide palette of chemical toxins and virulence factors that target the ribosome and associated translation factors (Bérdy, 2005), we reasoned that a screen of diverse bacterial strains could detect counter-surveillance drugs or virulence produced by bacteria. We screened bacterial species by growth of individual bacterial strains with a *C. elegans* strain carrying a mutation in a translation factor gene for bacterial activities that disrupt the induction of xenobiotic detoxification genes. This screen identified an activity of *Kocuria rhizophila* that suppresses *C. elegans* surveillance of translation (Govindan et al., 2015). Here we identify by genetic analysis of *K. rhizophila* that C_50_ carotenoids are the bacterial activity that suppresses the *C. elegans* translational toxin defense response. This *K. rhizophila* activity suppresses the induction of xenobiotic detoxification and food aversion responses caused by deficits in protein synthesis. Another bacterial species that also produces the C_50_ carotenoid decaprenoxanthin, *Corynebacterium glutamicum,* also suppressed the induction of xenobiotic detoxification genes in a *C. elegans* translation factor mutant, and *C. glutamicum* carotenoid biosynthesis mutants also fail to suppress *C. elegans* detoxification induction. The bacterial inhibition of this detoxification response increases the potency of bacterial toxins that target translation: treatment of wild type *C. elegans* with purified *K. rhizophila* C_50_ carotenoids caused hypersensitivity to drugs that inhibit translation. Thus, the evolution and lateral transfer between bacterial strains of these carotenoids may have been selected because they increase in the potency of bacterial toxins. Bile acid signaling mediates the signaling between *C. elegans* translational surveillance and induction of drug detoxification genes (Govindan et al., 2015). Addition of mammalian bile acids to the *K. rhizophila* inhibited *pgp-5* response pathway bypassed the carotenoid inhibition of the response, showing that carotenoids act upstream of bile acid production.

## Results

### Bacterial carotenoids suppress *C. elegans* surveillance and response to translational deficits

Like many animal species, *C. elegans* encodes about 500 xenobiotic detoxification genes ---for example, cytochrome p450, ABC transporter, UDP-glycosyl transferase genes--that are induced by particular toxins or genetic deficits (Melo and Ruvkun, 2012). Particular suites of *C. elegans* detoxification genes are strongly induced by inhibition of ribosomal proteins, tRNA synthetases, and other genes implicated in translation (Govindan et al., 2015). The *C. elegans* ABC transporter gene *pgp-5* is strongly induced by disruption of ribosomal and other translation factor genes but not by defects in other conserved cellular components such as the mitochondrion or the cytoskeleton (Govindan et al., 2015). A *pgp-5::gfp* fusion gene is therefore a reporter of *C. elegans* translational dysfunction and not just general poor health or poor growth conditions (Govindan et al., 2015). This ABC transporter protein may eliminate toxins that target the ribosome from the cell, but this has not been established. The translational defect can even be limited to the germline using the *C. elegans eft-3(q145)* germline-translation defective mutant, which is sterile, but with normal translation in somatic cells. In the *eft-3(q145)* mutant, *pgp-5p::gfp* is strongly induced in the intestine when the animals are fed the benign *E. coli* OP50 (Figure S1A; Figure 1A). Feeding *K. rhizophila* rather than *E. coli* to *C. elegans eft-3(q145); pgp-5p::gfp* animals disrupts the induction of *pgp-5p::gfp* (Figure S1A; Figure 1A-B) (Govindan et al., 2015). *K. rhizophila* is an Actinobacteria, a clade rich in drug biosynthetic pathways. *K. rhizophila* species are found in soils and rotting fruits associated with natural populations of *C. elegans* in orchards (Samuel et al., 2016). *K. rhizophila* species are also normal inhabitants of skin and mucous membranes of humans and animals, but can be associated with human infections (Becker et al., 2008) (Moissenet et al., 2012) (Purty et al., 2013).

**Figure 1:**
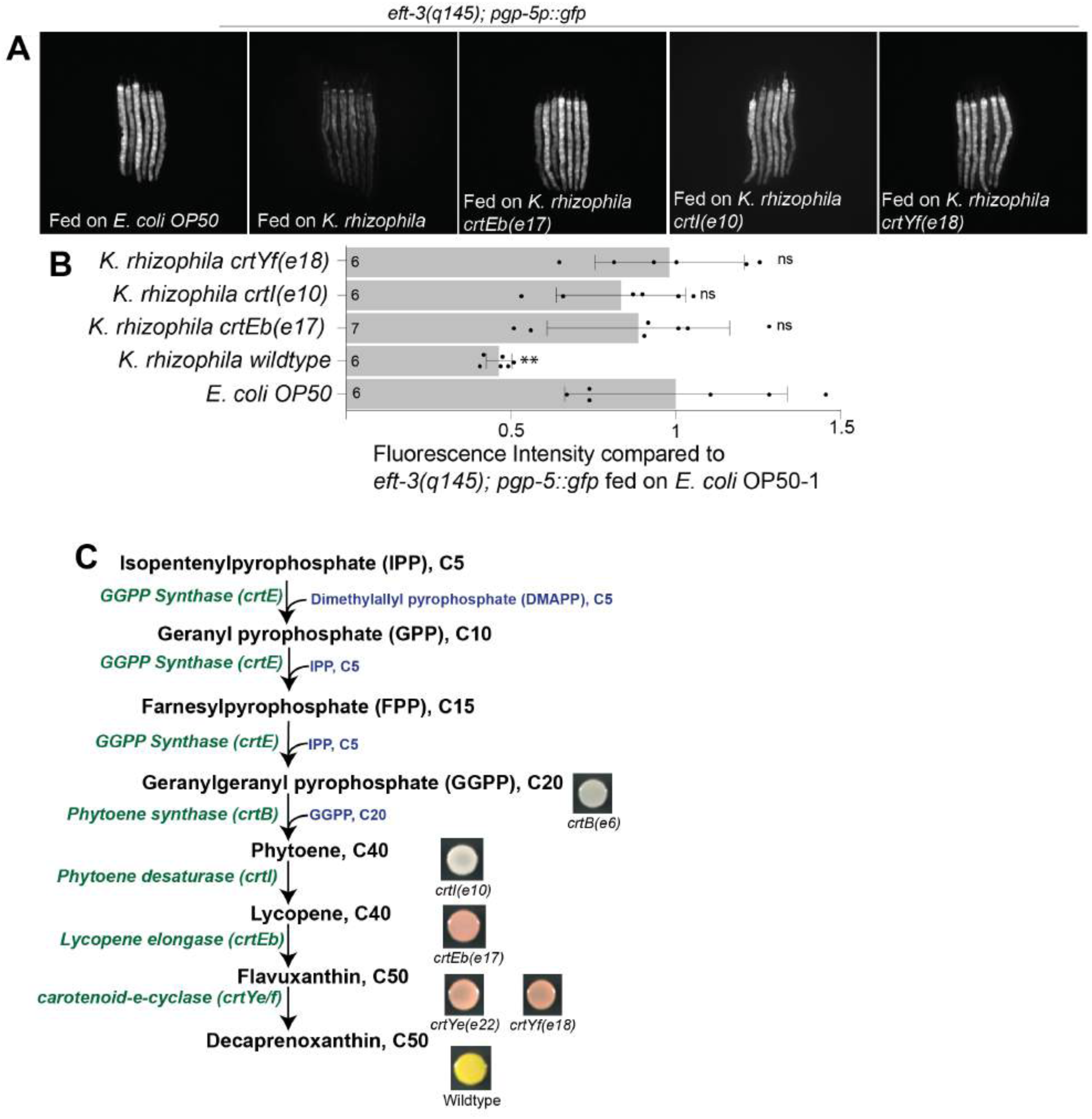
*K.rhizophila* carotenoid biosynthetic mutants fail to suppress *C. elegans* surveillance of translational deficits. A) *pgp-5p::gfp* induction was significantly reduced in *eft-3(q145); pgp-5p::gfp* animals cultivated on *K. rhizophila* wildtype while mutants in *K. rhizophila crtEb(e17)*, *K. rhizophila crtI(e10)*, and *K. rhizophila crtYe(e2)* did not suppress the GFP induction. B) *K. rhizophila* feeding reduced *pgp-5p::gfp* induction in *eft-3(q145); pgp-5p::gfp* animals significantly, while in the *K. rhizophila* mutants, *pgp-5p::gfp* expression in this strain was not affected. Unpaired t-test, **P<0.01. Mean ± s.d is shown. The number of animals analyzed per condition is shown above each bar. ns, was not significant compared to *eft-3(q145); pgp-5p::gfp* fed on *E. coli* OP50 C) The *Kocuria rhizophila* C_50_ carotenoid biosynthetic pathway. The genome of *K. rhizophila* contains an operon that encodes predicted carotenoid (*crt*) biosynthetic genes: *crtE* (KRH_20850; encoding GGPP synthase), *crtB* (KRH_20840; encoding phytoene synthase), *crtI* (KRH_20830; encoding phytoene desaturase), *crtEb* (KRH_20800; encoding lycopene elongase), *crtYe* (KRH_20820; encoding C_50_ carotenoid epsilon cyclase subunit) and *crtYf* (KRH_20810; encoding C_50_ carotenoid epsilon cyclase subunit). The crtYe and crtYf subunits form a heterodimeric cyclase complex that catalyzes generation of C_50_ carotenoid. Colonies of wildtype K. rhizophila and mutants are shown. While wildtype *K. rhizophila* colonies were yellow, *crtB* and *crtI* mutant colonies were colorless. *crtYe*, *crtYf*, and *crtEb* mutant colonies were various shades of red.

To establish that *K. rhizophila* can suppress the surveillance of a range of defects in translation, we tested the induction of the *pgp-5p::gfp* xenobiotic detoxification gene reporter after gene inactivation by RNAi of other genes that encode ribosomal protein subunits or translation factors. For example, growth of wild type *C. elegans* carrying *pgp-5p::gfp* on *E. coli* engineered to produce *rpl-1* dsRNA or *vrs-2* dsRNA, which inactivate the RPL-1 ribosomal protein or the VRS-2 tRNA synthetase genes, caused induction of *pgp-5p::gfp.* By contrast, in animals fed on either *rpl-1* dsRNA or *vrs-2* dsRNA and transferred to *K. rhizophila*, *pgp-5p::gfp* expression was decreased (Figure S1B-C). The suppression of surveillance pathways by *K. rhizophila* is specific for translational defects because *K. rhizophila* does not suppress the induction of the mitochondrial stress response or endoplasmic reticulum stress response (Govindan et al., 2015). Constant exposure to *K. rhizophila* is necessary to suppress *C. elegans* translational surveillance: *eft-3(q145);pgp-5p::gfp* animals fed on *K. rhizophila* and transferred after various times to *E. coli* OP50 plates re-expressed *pgp-5p::gfp* within 12 hours of transfer (Figure S1D). To identify the *K. rhizophila* genetic pathways responsible for the inhibition of *pgp-5p::gfp* induction by a *C. elegans* deficit in translation, we conducted a forward genetic screen after EMS mutagenesis of *K. rhizophila* for bacterial mutant strains that can no longer inhibit *pgp-5p::gfp* induction in *eft-3(q145)*. ∼2000 individual *K. rhizophila* strains that grew normally on bacterial LB plates after EMS mutagenesis were fed to *C. elegans eft-3(q145); pgp-5p::gfp* animals, one bacterial mutant per well, and screened for *K. rhizophila* mutants that failed to suppress *C. elegans pgp-5p::gfp* induction in a population of 50-100 *eft-3(q145); pgp-5p::gfp* animals. We identified six *K. rhizophila* mutant strains that failed to inhibit the induction of *pgp-5p::gfp* in the *eft-3(q145); pgp-5p::gfp* strain (Figure S1E-F). All these *Kocuria* mutant strains had defects in colony pigmentation compared to wild type *K. rhizophila* (Figure S2A). Wildtype *K. rhizophila* is yellow, whereas the six mutant colonies were red or white or orange. Using this discoloration phenotype, we visually screened ∼500,000 bacterial colonies after EMS mutagenesis for pigmentation mutants; 71 mutants were identified (Figure S2B). The 71 mutants along with 25 control non-discolored mutants were tested on *C. elegans eft-3(q145); pgp-5p::gfp* animals and scored for GFP induction. All 71 pigmentation mutants failed to suppress *pgp-5p::gfp* induction in the *C. elegans eft-3(q145)* mutant, whereas the 25 normally colored *K. rhizophila* strains suppressed *pgp-5p::gfp* induction, like wild type *K. rhizophila* (Table S1; Figure S2C; Figure 1A-B).

Genome sequencing of 23 of these pigmentation mutants revealed that each carried a mutation in one of six carotenoid biosynthetic cluster genes (Figure S2D-E). Carotenoids are yellow to red colored pigments, which are produced by a terpenoid biosynthetic pathway (Takarada et al., 2008). The reaction catalyzed by GGPP synthase CrtE, phytoene synthase CrtB and phytoene desaturase CrtI mediate steps in the production of lycopene (Klassen, 2010) (Krubasik et al., 2001a). CrtEb and CrtYe/f cyclases catalyze the biosynthesis of C_50_ carotenoid from lycopene. C_50_ carotenoids are synthesized more rarely than other carotenoids (Krubasik et al., 2001a). Six missense mutations (*e21*, *e11*, *e23*, *e14*, *e5*, and *e13*) and two nonsense mutations (*e15* and *e10*) were in *crtI*, which encodes phytoene desaturase (Figure 1C; Figure S2D-E). CrtI catalyzes the conversion of the non-colored phytoene to lycopene, which is red. All these *crtI* mutants are white colored colonies (Figure S2D; Figure 1C) similar to the *Corynebacterium glutamicum* D*crtI* mutant (Heider et al., 2012) (Figure S2F) and thus are predicted to be defective in lycopene synthesis. The six missense mutations are in highly conserved residues suggesting that these are strong loss of function mutations (Figure S3). The *e4*, *e6*, and *e8* are missense mutations in the *crtB* gene, which encodes phytoene synthase (Figure 1C; Figure S2D-E). These missense mutations are in highly conserved residues, suggesting that these are strong loss of function mutations (Figure S4). The mutants produce white bacterial colonies (Figure S2D; Figure 1C), as was observed in the *C. glutamicum* D*crtB* mutant (Heider et al., 2012) (Figure S2F). We obtained four mutations in *crtEb*; two nonsense mutations (*e16* and *e17*) and two missense mutations (*e3* and *e19*) in highly conserved residues (Figure 1C; Figure S2E; Figure S5). Mutations in *crtEb* are likely to be defective in the conversion of lycopene to flavuxanthin (Figure 1C). These mutants form pale red colonies (Supplementary Fig. 2d; Fig. 1c) probably because of accumulation of lycopene (but not flavuxanthin), as is seen in the *C. glutamicum* D*crtEb* mutant (Heider et al., 2012) (Figure S2F). *e17* is an early stop mutation in *crtEb* which is predicted to produce a truncated protein of 13 amino acids (Figure S5). Mutations in *crtYe* and *crtYf* are defective in the last step; the homologs of these genes in *C. glutamicum* are known to catalyze synthesis of decaprenoxanthin (Krubasik et al., 2001a) (Monnet et al., 2010). Mutations in *crtYe* and *crtYf* produce pale red to orange colonies (Figure S2D). A *C. glutamicum* D*crtY* mutant that deletes both *crtYe* and *crtYf* accumulates flavuxanthin and is a pale orange to red color (Heider et al., 2012) (Figure S2F). Even though several bacterial species contain the CrtEb and CrtYe/f genes (Figure S6; See Supplementary methods), the only genetically and biochemically well-characterized C_50_ carotenoid is decaprenoxanthin from *C. glutamicum* (Heider et al., 2012). In *C. glutamicum*, the enzymes CrtEb, CrtYe, and CrtYf convert lycopene to the C_50_ carotenoid decaprenoxanthin (Krubasik et al., 2001a) (Monnet et al., 2010) (Figure 1C). CrtEb catalyzes the elongation of C_40_ acyclic lycopene to acyclic C_50_ carotenoid flavuxanthin (Krubasik et al., 2001a). CrtYe and CrtYf form a C_50_ cyclase which converts C_50_ flavuxanthin to decaprenoxanthin (Krubasik et al., 2001a) (Krubasik et al., 2001b). *K. rhizophila crtYe* and *crtYf* are 38% and 34% identical to *C. glutamicum crtYe* and *crtYf* respectively. Therefore, the yellow pigment produced by *K. rhizophila* may belong to the decaprenoxanthin C_50_-subfamily.

Consistent with the anti-*C. elegans* translational surveillance activity of *K. rhizophila* decaprenoxanthin, we found that *C. glutamicum* ATCC13032 which produces a similar C_50_ pigment also suppressed *pgp-5p::gfp* induction in *eft-3(q145);pgp-5p::gfp* (Figure 2A-B). And like the *K. rhizophila* carotenoid mutants, *C. glutamicum* ATCC13032 deletion mutants in crtY, crtEb, crtI, and, crtB which disrupt decaprenoxanthin production (Heider et al., 2012) allowed the induction of *pgp-5p::gfp* in *eft-3(q145);pgp-5p::gfp* animals (Figure 2A-B). Thus, the *C. glutamicum* carotenoid mutations also fail to suppress *pgp-5p::gfp* induction in *eft-3(q145)*.

**Figure 2:**
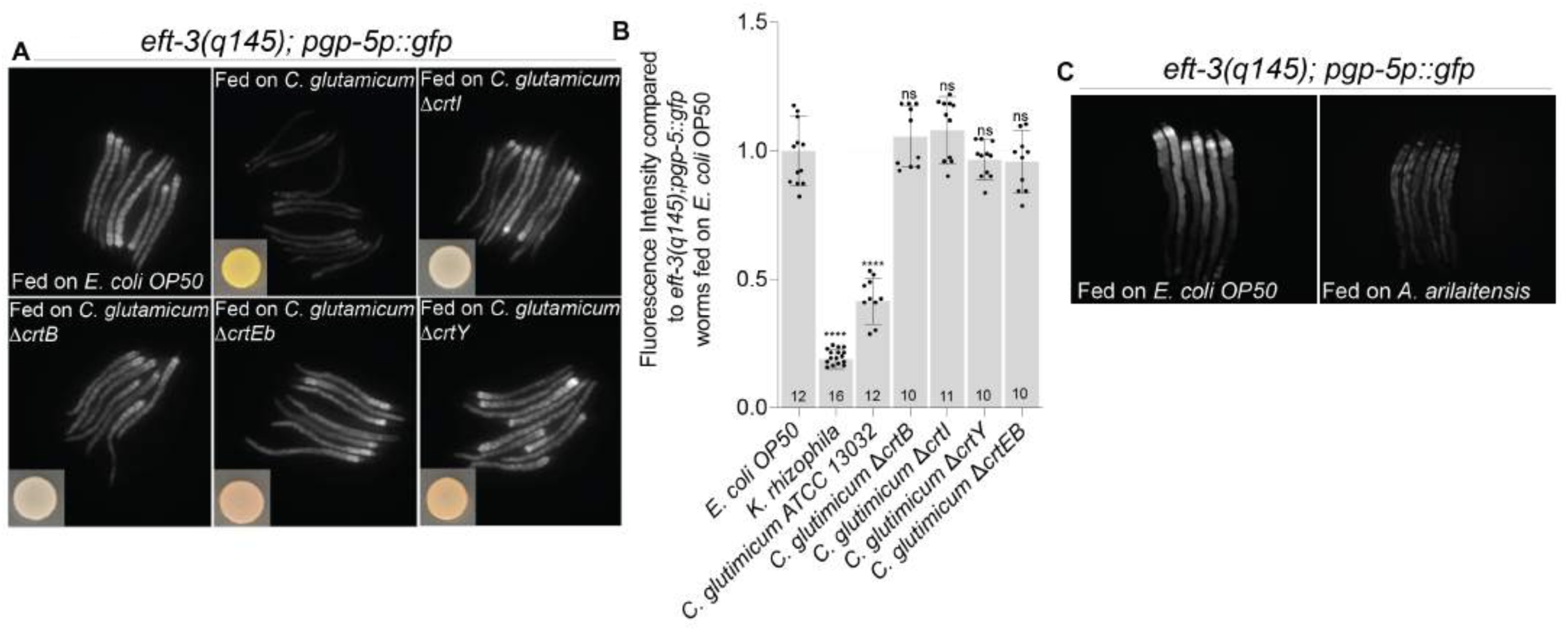
Bacterial C50 carotenoids suppress *C. elegans* surveillance of translational deficits. A) *pgp-5p::gfp* induction was significantly reduced in *eft-3(q145); pgp-5p::gfp* animals grown on *C. glutamicum* wildtype while mutants in *C. glutamicum* D*crtEb*, D*crtI*, D*crtY*, and D*crtB* did not suppress the GFP induction. B) Quantification of *pgp-5p::gfp* expression in *eft-3(q145); pgp-5p::gfp* animals grown on *C. glutamicum* wildtype, D*crtEb*, D*crtI*, D*crtY*, and D*crtB* mutants. Unpaired t-test, ****P<0.0001. Mean ± s.d is shown. The number of animals analyzed per condition is shown above each bar. ns, was not significant compared to *eft-3(q145); pgp-5p::gfp* fed on *E. coli* OP50. C) *pgp-5p::gfp* induction was significantly reduced in *eft-3(q145); pgp-5p::gfp* animals grown on wildtype *A. arilaitensis*.

*Arthrobacter arilaitensis,* the source of the yellow pigment in Livarot, Maroilles, Munster, Limburger and Tilsit cheeses, also produces decaprenoxanthin (Monnet et al., 2010). Feeding *A. arilaitensis* to *eft-3(q145); pgp-5p::gfp* animals also suppresses *pgp-5p::gfp* induction ((Figure 2C). Thus, *C. glutamicum, A. arilaitensis,* or *K. rhizophila* produce a pigmented carotenoid that suppresses translational surveillance, and mutations in *C. glutamicum* or *K. rhizophila* that disrupt decaprenoxanthin biosynthesis inactivate this suppression of surveillance.

One trivial explanation for the failure of the *K. rhizophila* carotenoid mutants to suppress *pgp-5p::gfp* in a translation-defective *C. elegans* mutant would be that these *K. rhizophila* pigmentation mutants might induce *pgp-5p::gfp* even in a wildtype *C. elegans* background. But wild type *C. elegans* carrying *pgp-5p::gfp* grown on the *K. rhizophila crtEb(e17)*, *crtYe(e22)*, *crtYf(e18)*, *crtB(e6)* or *crtI(e10)* mutants do not induce *pgp-5p::gfp* (Table S1; Figure S7A). Another possible interpretation was that *K. rhizophila* feeding might induce other stress responses in *C. elegans* that somehow “distract” the animal from surveillance of translation. We tested the induction of other GFP fusion reporters of stress on wild type and various *K. rhizophila* mutants, and found that *K. rhizophila* wildtype or carotenoid mutants did not induce *hsp-4p::gfp* ((Figure S7B), *hsp-6p::gfp* (Figure S7C), F35E12.5p::gfp (Figure S7D), or *clec-60::gfp* (Figure S7E). These stress reporter experiments rule out trivial explanations, such as that *Kocuria* itself induces the expression of *pgp-5::GFP,* or that *Kocuria* induces a distracting stress response so that translation deficits are no longer coupled to *pgp-5::GFP* induction. These experiments also rule out that the carotenoid pigment of wild type *Kocuria* absorbs the UV light used to excite GFP fusion genes or the green light emitted by GFP fusion proteins.

### Purified carotenoids from *Kocuria rhizophila* suppress *C. elegans* translational surveillance

Carotenoids such as decaprenoxanthin are lipophilic molecules that localize to the cell membrane and can be extracted in non-polar solvents. We extracted wild type *K. rhizophila* cultures with methanol (Figure S8; Figure 3A-C). TLC analysis of the extract revealed the presence of yellow-orange pigments (Figure 3A, inset). High resolution LC-MS analysis of a methanol extract from *K. rhizophila* showed a complex mixture containing at least six components with similar absorbance spectra that eluted from a reverse phase C18 column (Figure 3A-B; See supplemental methods section). Average absorbance spectra of the elution peaks revealed absorption maxima at 414, 438 and 468 nm, which is similar to the published absorption spectra of decaprenoxanthin from *Arthrobacter* (Giuffrida et al., 2016) (Sutthiwong and Dufossé, 2014). For three of the elution peaks at 19, 16, and 9 min, high resolution mass spectra contained ions that were consistent with decaprenoxanthin and its mono- and di-glucosides, respectively, with mass accuracy within 2 ppm (Figure 3C). Three other peaks with similar absorbance spectra were not assignable by mass.

**Figure 3:**
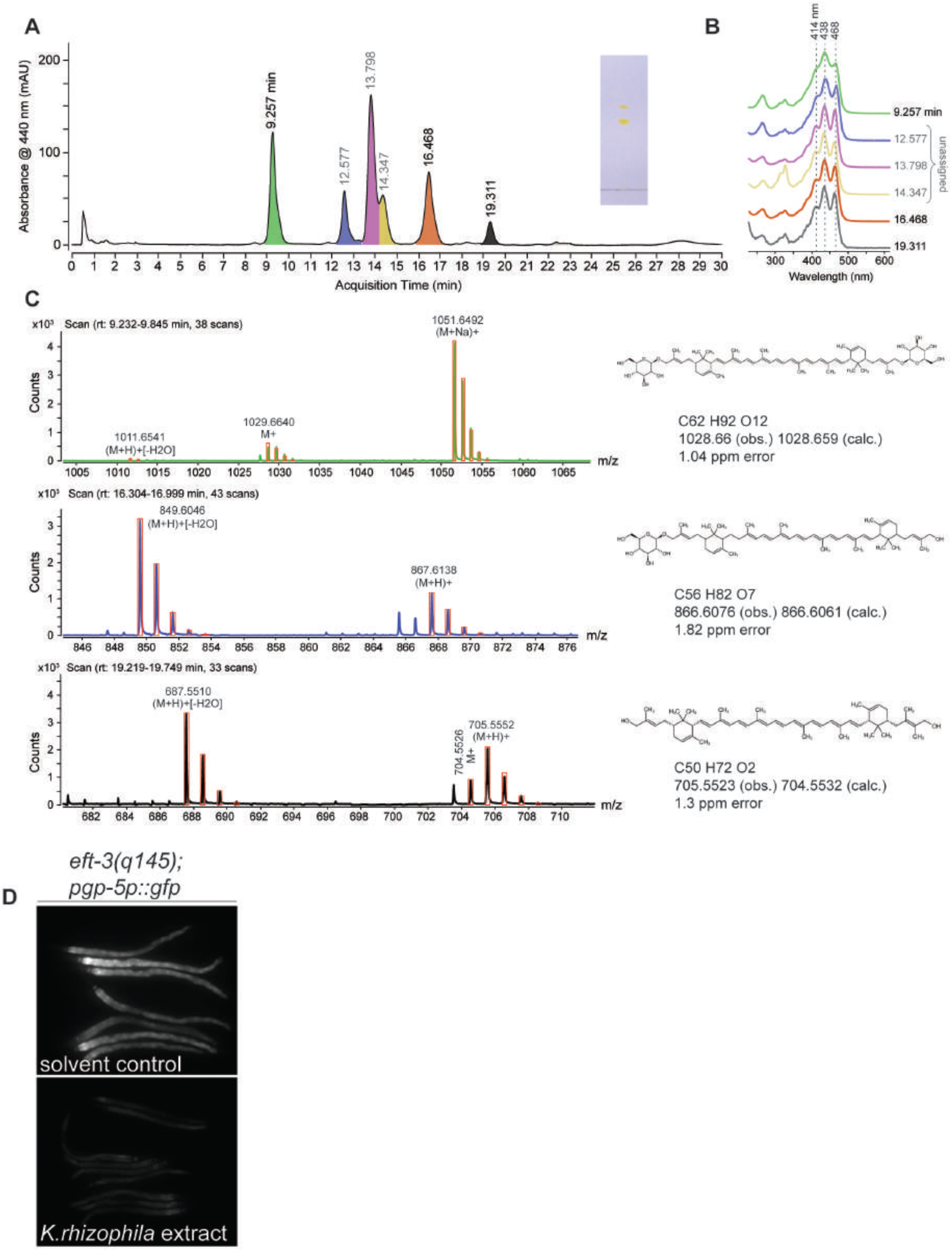
LC-MS analysis of carotenoid extracts from *K. rhizophila*. A) Chromatogram of 440 nm absorbance after reverse phase LC separation of clarified extracts on a C18 column. Six major elution peaks are indicated, Inset: TLC of *K. rhizophila* extract showing yellow-orange pigments. B) Average absorbance spectra for each indicated elution peak. Spectra are normalized and baseline shifted for clarity. All six elution peaks share similar absorbance spectra, but three peaks (grey lettering) were not conclusively assigned by mass. C) Background-subtracted averaged MS spectra of three assigned elution peaks at approximately 9 (green), 16 (orange), and 19 min (dark grey) with observed neutral monoisotopic mass (obs.), corresponding exact calculated mass (calc.), and mass error for each indicated chemical formula. Each spectrum shows the calculated isotopic distribution for the identified adduct ions (red bars). D) 750µg/ml of *K. rhizophila* extract inhibited *pgp-5p::gfp* induction in *eft-3(q145); pgp-5p::gfp* animals.

To analyze the carotenoid production in the *K. rhizophila* mutants, we extracted carotenoids from *crtI(e10)*, *crtEb(e17)*, *crtYe(e22)*, and *crtYf(e18)*, which are nonsense mutant alleles and *crtB(e6)*, a missense mutant. Spectrophotometric analysis of methanol extracts from wildtype *K. rhizophila* showed absorption maxima of 415-425nm, whereas the *crtEb(e17)* extract showed absorption maxima at 445-455nm (Figure S9). *K. rhizophila crtEb(e17)* and *crtYe(e22)* mutant methanol extract show similar absorption spectra while the extract from *K. rhizophila crtI(e10)* and *crtb(e6)* showed no absorption at all. The methanol extracts from *crtYf(e18)* showed two separate absorption peaks one at ∼400 nm and another minor one at ∼500nm. To determine the identities of these absorption peaks in the mutants, we conducted LC-MS analysis of carotenoid extracts from *crtYe(e22)* and *crtEb(e17)* (Figure S10A-C). Mutations in *crtEb* are predicted to block the synthesis of carotenoids at the flavuxanthin biosynthesis step and thus are likely to accumulate lycopene (Figure S10C). Consistent with this prediction, MS spectral analysis revealed that *crtEb(e17)* mutants accumulate lycopene. Mutations in *crtYe* are predicted to block the synthesis of carotenoids at the decaprenoxanthin biosynthesis step and thus likely to accumulate flavuxanthin (Figure S10C). However, we found that *crtYe(e22)* mutants accumulate lycopene as well as other unidentified peaks.

We tested whether the crude methanol extracts from wildtype *K. rhizophila,* containing pigmented carotenoids, could suppress GFP induction in the germline-translation defective *C. elegans eft-3(q145);pgp-5p::gfp* animals. Animals fed on *E. coli* with the *K. rhizophila* carotenoid extract exhibited significantly reduced *pgp-5p::gfp* expression compared to *eft-3(q145);pgp-5p::gfp* animals fed on *E. coli* with control methanol extract (Figure 3D). The wild type *K. rhizophila* carotenoid methanol extract could also rescue the failure to suppress *C. elegans* surveillance by *K. rhizophila* carotenoid biosynthetic mutants: when *eft-3(q145);pgp-5p::gfp* animals were fed on *K. rhizophila crtEb(e17)*, *crtB(e6)* or *crtI(e10)* mutants supplemented with wildtype *K. rhizophila* methanol extract, GFP was not induced while in the animals fed on control extract, the GFP expression was induced by the *C. elegans eft-3* mutation (Figure 4A).

**Figure 4:**
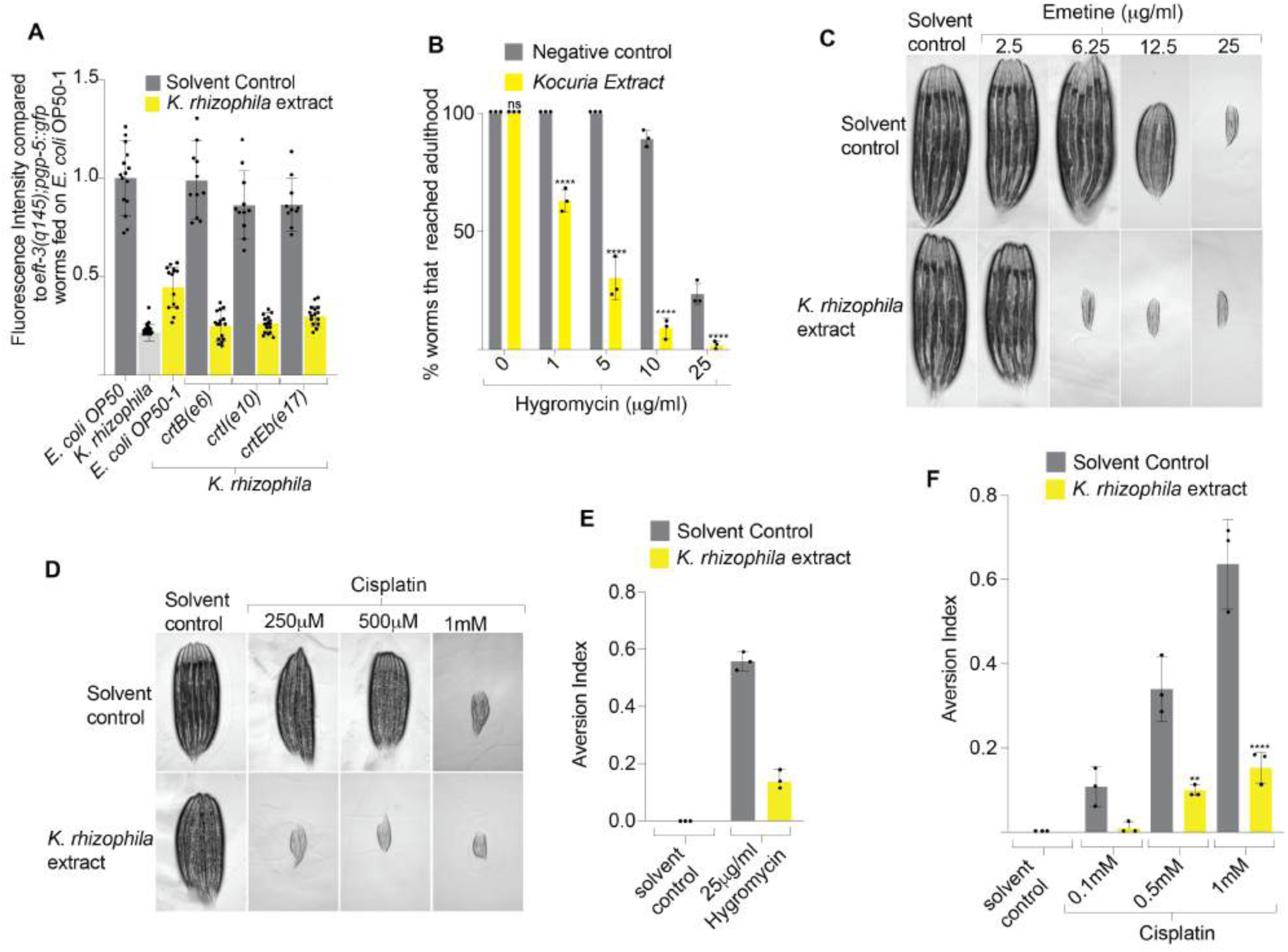
Purified carotenoids from *Kocuria rhizophila* suppress *C. elegans* translational surveillance. A) Quantification of *pgp-5p::gfp* expression in *eft-3(q145); pgp-5p::gfp* animals fed on *K. rhizophila* wildtype, *K. rhizophila crtEb(e17)*, *K. rhizophila crtI(e10)*, and *K. rhizophila crtEb(e6)* supplemented with either control extract or an extract from wild type *K. rhizophila*. B) Animals treated with *K. rhizophila* carotenoid extract were hypersensitive to the translational inhibitor hygromycin. Unpaired t-test, ****P<0.0001 compared to the wildtype worms fed on *E. coli* OP50 containing solvent extract and hygromycin. Mean ± s.d is shown. Data were collected from three independent trials of at least 20 animals for each condition. ns, was not significant compared to wildtype grown on *E. coli* OP50 containing solvent extract with no hygromycin C) Animals treated with *K. rhizophila* carotenoid extract were hypersensitive to emetine. D) Animals treated with *K. rhizophila* carotenoid extract were hypersensitive to cisplatin. E) Animals treated with *K. rhizophila* carotenoid extract failed to show aversive behaviors on hygromycin compared animals treated with control solvent and hygromycin. F) Animals treated with *K. rhizophila* carotenoid extract failed to avoid cisplatin-compared animals treated with control solvent and cisplatin. Unpaired t-test, ****P<0.01, **P<0.0001. Mean ± s.d is shown. ns, not significant compared to

Because C_50_ carotenoids have multiple conjugated double bonds, they are likely to be antioxidants (Edge et al., 1997). Several carotenoids including C_30_, C_40_, or C_50_ carotenoids are antioxidants. However, it is unlikely that the ROS-quenching property of carotenoids is responsible for the suppression of *pgp-5p::gfp* induction. First, multiple antioxidants do not suppress the induction of *pgp-5p::gfp* in a *C. elegans* translation defective mutant: *eft-3(q145);pgp-5p::gfp* animals grown on *E. coli* OP50 with N-acetyl cysteine, ascorbic acid, Trolox, or resveratrol induce *pgp-5p::gfp* normally (Figure S11A). Second, *pgp-5p::gfp* is not induced by oxidative stress (Govindan et al., 2015).

Consistent with the genetic analysis of *K. rhizophila* and *C. glutamicum* highlighting the production of C_50_ decaprenoxanthin as key to suppression of *C. elegans* immune responses, we found that other commercially available 40 carbon carotenoids in the pathway to decaprenoxanthin C_50_ biosynthesis cannot suppress the induction of *pgp-5p::gfp* in a *C. elegans* translation defective mutant. For example, treating *eft-3(q145);pgp-5p::gfp* animals grown on *E. coli* OP50 with either C_40_ beta-carotene or C_40_ astaxanthin did not suppress *pgp-5p::gfp* induction (Figure S11B). C_40_ zeaxanthin, C_40_ neurosporene, C_40_ violaxanthin, C_40_ delta-carotene, or C_40_ alpha-carotene also did not suppress *pgp-5p::gfp* induction (Figure S11C). These C_40_ carotenoids cannot suppress the response to translational deficits like the C_50_ carotenoids produced by *K. rhizophila* and *C. glutamicum*.

### Suppression of *C. elegans* translational surveillance by *Kocuria rhizophila* carotenoids enhances the toxicity of ribosomal toxins

*K. rhizophila* also suppresses the *C. elegans* detoxification response to translation inhibiting drugs. Hygromycin is a bacterially produced antibiotic (from *Streptomyces hygroscopicus*) that inhibits translation and induces xenobiotic detoxification in *C. elegans*. While 10 µg/ml of hygromycin induces *pgp-5p::gfp* expression in wild type animals grown on *E. coli* OP50, wild type animals grown on *K. rhizophila* and 10µg/ml of hygromycin fail to induce *pgp-5p::gfp* (Figure S7A; Figure S11D). However, at high concentrations of hygromycin, *pgp-5p::gfp* is induced both in animals fed on *E. coli* OP50 or *K. rhizophila* (Figure S11D). By contrast, wild type *C. elegans* carrying *pgp-5p::gfp* grown on the *K. rhizophila* carotenoid biosynthesis mutants *crtI(e10)* or *crtEb(e17)* induced GFP at the low concentration of 10µg/ml of hygromycin unlike animals grown on wild type *K. rhizophila* that produces the carotenoid (Figure S7A). Similar results were obtained with emetine which blocks protein synthesis by binding to the 40S subunit of the ribosome; wild type *K. rhizophila* suppressed the *pgp-5p::gfp* induction by a low dose of emetine but the carotenoid mutant *K. rhizophila* now allowed induction of *pgp-5p::gfp* by a low dose of emetine (Figure S12A-B).

We tested whether the suppression of drug detoxification responses by *K. rhizophila* carotenoids increase *C. elegans* sensitivity to translational inhibitors. While wildtype animals grown on 10µg/ml hygromycin when fed *E. coli* OP50 are not growth inhibited, animals grown on 10µg/ml hygromycin plus *K. rhizophila* carotenoid extract when fed *E. coli* OP50 are strongly growth inhibited (Figure 4B; Figure S12C). Carotenoids without hygromycin are not toxic to the worms (Figure 4B; Figure S12C). Similar results were obtained with emetine: animals treated with *K. rhizophila* extracts were hypersensitive to emetine compared to animals fed on control extract (Figure 4C). The xenobiotic hypersensitivity phenotype is specific to translation defects because *K. rhizophila* carotenoid extract does not alter the sensitivity of animals to antimycin, a mitochondrial poison (Figure S12E).

*pgp-5p::gfp* is also induced in response to genotoxic stress: cisplatin, which interferes with DNA replication, induces *pgp-5p::gfp*. While 1 mM cisplatin induces *pgp-5p::gfp* expression in wild type *C. elegans* grown on *E. coli* OP50, wild type *C. elegans* grown on *K. rhizophila* and 1mM cisplatin fail to induce *pgp-5p::gfp* (Figure S12D). By contrast, *pgp-5p::gfp* animals fed *K. rhizophila crtI(e10), crtEb(e17)*, or *crtYe(e22)* carotenoid mutant bacteria induce *pgp-5p::gfp* in response to 1 mM cisplatin normally (Figure S12D). And the genotoxic response is salient for survival of cisplatin: animals treated with *K. rhizophila* extracts were hypersensitive to cisplatin compared to animals fed on control extract (Figure 4D).

*C. elegans* food aversion behaviors are induced when animals are exposed to xenobiotics or essential gene inactivations (Melo and Ruvkun, 2012). Exposing animals to hygromycin or cisplatin induces strong food aversion, and this food aversion is modulated by *K. rhizophila* carotenoid extract (Figure 4E-F). While ∼40% of animals exposed to 25µg/ml hygromycin display food aversion behavior, only ∼15% of animals exposed to 25µg/ml hygromycin and *K. rhizophila* carotenoid extract show aversion (Figure 4E). Similar results were obtained with cisplatin: ∼50% of animals exposed to 1mM cisplatin display aversion behavior while <20% of animals exposed to 1mM cisplatin and *K. rhizophila* carotenoid extract display food aversion (Figure 4F).

### *C. elegans* pathway analysis of *K. rhizophila* inhibition of translational surveillance

How do *K. rhizophila* carotenoids inhibit the induction of xenobiotic detoxification response pathways? To address this question, we conducted genetic epistasis analysis with *C. eleg*ans mutations that disrupt or activate the signal transduction pathway for translational surveillance at various steps (Govindan et al., 2015). The *zip-2*/bZIP transcription factor is required for the induction of *pgp-5p::gfp* expression in response to translation inhibition (Govindan et al., 2015). Constitutive expression of ZIP-2::mCherry from an intestine-specific promoter fusion is sufficient to induce *pgp-5p::gfp* expression in wildtype *C. eleg*ans without translation inhibition (Fig. 5a,b). *pgp-5p::gfp* induction in this ZIP-2::mCherry constitutive expression strain was similar in animals fed on *E. coli* OP50 or wild type *K. rhizophila* (Figure A-B), suggesting that the *K. rhizophila* carotenoids disrupt a surveillance pathway component upstream of the production of the ZIP-2 bZIP transcription factor.

**Figure 5:**
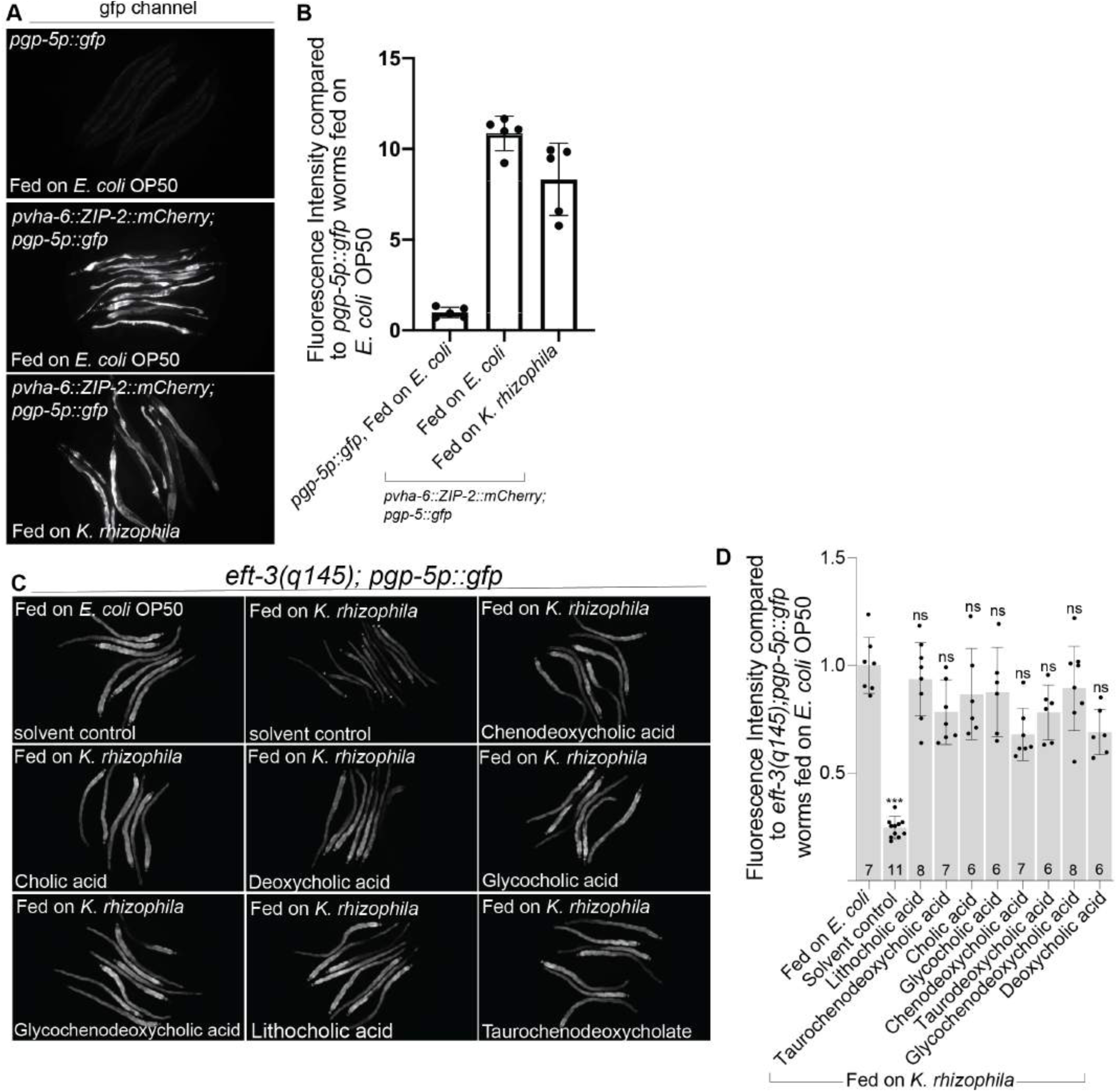
*C. elegans* pathway analysis of *K. rhizophila* inhibition of translational surveillance. A) *pgp-5p::gfp* was constitutively induced in animals fed on *E. coli* OP50 or *K. rhizophila* expressing ZIP-2::mCherry in the intestine under the control of the *vha-6* promoter. B) Quantification of *pgp-5p::gfp* activation in animals expressing ZIP-2::mCherry in the intestine under the control of *vha-6* promoter. Compared to the *pgp-5p::gfp* animals fed on *E. coli* OP50, animals expressing ZIP-2::mCherry in the intestine under the control of *vha-6* promoter had increased *pgp-5p::gfp* levels when fed on either *E. coli* OP50 or *K. rhizophila*. Mean ± s.d is shown. The black dots on each bar indicates each animal analyzed per condition. C) Supplementation of bile acids suppressed the *K. rhizophila pgp-5p::gfp* activation defect in *eft-3(q145); pgp-5p::gfp* animals. D) Quantification of the suppression of the *K. rhizophila pgp-5p::gfp* activation defect in *eft-3(q145); pgp-5p::gfp* animals by bile acids shown in Figure 4B. Unpaired t-test, ***P<0.0001. Mean ± s.d is shown. The number of animals analyzed per condition is shown above each bar. ns, was not significant compared to *eft-3(q145); pgp-5p::gfp* fed on *E. coli* OP50.

The induction of *C. elegans* xenobiotic detoxification genes by translation inhibition depends on multiple steps in the bile acid biosynthetic pathway from cholesterol (Govindan et al., 2015). *C. elegans* with defects in bile acid biosynthesis fail to activate *pgp-5p::gfp* in response to inhibition of translation. Although the *C. elegans* bile acid biosynthetic pathway is necessary for *pgp-5p::gfp* in response to inhibition of translation, bile acids are not sufficient to induce *pgp-5p::gfp* in wildtype animals (Govindan et al., 2015). But addition of mammalian bile acids to these *C. elegans* bile acid pathway mutants substitutes for the missing endogenous bile acids to reanimate this translational defect signal (Govindan et al., 2015). While *K. rhizophila* feeding inhibits the induction of *pgp-5p::gfp* in *eft-3(q145)* animals, addition of exogenous mammalian bile acids reactivates GFP expression when feeding on wild type *K. rhizophila* (Figure 5C-D). Thus, *K. rhizophila* carotenoids act either upstream or at the bile acid signaling step of this *C. elegans* translational surveillance and response pathway.

## Discussion

We have found that multiple actinobacterial species produce C_50_ carotenoid pigments that are potent inhibitors of the animal surveillance pathways for bacterial toxins and virulence factors which target translation. We genetically dissected how the actinobacteria *Kocuria rhizophila* inhibits the surveillance and detoxification response of *C. elegans* to translational deficits. Our bacterial genetic analysis identified the carotenoid biosynthetic pathway as the key bacterial countermeasure. We isolated mutants in the *K. rhizophila* C_50_ carotenoid biosynthetic pathway that failed to inhibit this *C. elegans* xenobiotic detoxification response (Figure 1A-C). Hydrophobic and pigmented extracts of *K. rhizophila* enriched for C_50_ carotenoid suppressed *C. elegans* xenobiotic detoxification response and restored the ability of *K. rhizophil*a carotenoid mutants to suppress the *C. elegans* response to defects in translation (Figure 2B-C). The *K. rhizophila* extracts also suppress *C. elegans* detoxification of toxins that target translation or DNA damage, rendering the animal far more sensitive to these toxins (Figure 4B-D, Figure S12C). Thus, carotenoid biosynthesis increases the potency of other bacterial toxins or virulence factors that might target translation or DNA.

A distantly related bacterium, *Corynebacterium glutamicum,* also suppresses this *C. elegans* xenobiotic detoxification response. Mutations that disable steps in the synthesis of this carotenoid no longer suppress *C. elegans* translational deficit responses--showing that C_50_ carotenoid regulation of the *C. elegans* surveillance is not limited to one particular strain of bacteria. C_50_ carotenoids suppress the induction of xenobiotic detoxification by inhibiting *C. elegans* bile acid signaling, which transduce translational deficits to defense responses (Govindan et al., 2015) (Figure 5C-D). The lipid solubility of the bacterial carotenoids and their abundance may be a key feature of their anti-bile acid signaling function in *C. elegans* surveillance of translational and DNA damage response.

The evolutionary rationale for the disabling of eukaryotic toxin response pathways by bacteria is that they could enhance the toxicity of ribosomal toxins. But neither wild type *K. rhizophila* nor the C_50_ biosynthesis mutants show evidence of translational inhibitor production (for example induction of *C. elegans pgp-5::gfp*). It is possible that *K. rhizophila* generates other toxins that cause deficits not related to translation, which are rendered less defended by the associated C_50_ production. Alternatively, *K. rhizophila* may associate with other bacteria that produce toxins or virulence factors that target the animal translational apparatus or cause DNA damage.

Carotenoids have been most studied in photosynthetic bacteria and plants, where they are auxillary light absorbing components in photosynthesis (Edge et al., 1997). Because carotenoids are antioxidants and light absorbing in photosynthetic bacteria and plants, the literature on carotenoids is highly focused on photosynthesis, light absorption, and free radical reactivity. In photosynthetic chlorophyll clusters, the carotenoids absorb and transduce light energy to photosynthetic electron transport systems to generate a pH gradient. In photosynthetic chlorophyll clusters, the carotenoids absorb and efficiently transduce light energy to photosynthetic electron transport systems. What is common between the *K. rhizophila* and *C. glutamicum* and photosynthetic bacteria is the strong absorbance in the visible light wavelengths and the lipid solubility of these pigments. These bacteria are highly colored because the carotenoids are both highly abundant and because they have conjugated double bond systems. Carotenoids are also antioxidants in photosynthesis. However, a simple antioxidant activity of carotenoids does not explain the suppression of the animal drug detoxification response because other antioxidants could not suppress *C. elegans* surveillance.

How do the C_50_ carotenoids of *K. rhizophila* and *C. glutamicum* inhibit animal surveillance of their translation competence? The simplest model is that C_50_ carotenoids induce a change in membrane fluidity to change cell signaling via bile acids. We found that the *K. rhizophila* C_50_ carotenoids suppress the induction of the *C. elegans* xenobiotic detoxification response by inhibiting a bile acid biosynthetic pathway. Although bile acids were traditionally thought to be emulsifiers of fat, thus aiding their enzymatic metabolism, bile acids are also signaling molecules in metabolism and immune pathways (Fiorucci et al., 2018). The lipid solubility of carotenoids may be a key feature of their anti-bile acid signaling function in *C. elegans* surveillance of translational and DNA damage response. Emulsification of lipids by bile acids may be relevant to the antagonistic relationship we show for bile acid signaling in *C. elegans* and the bacterial carotenoid inhibitors of that signaling pathway.

It is possible that the pigmentation of the C_50_ carotenoids, the particular wavelengths of light absorbance, are key to their anti-surveillance function. For example, our genetic analysis showed that the carotenoids of *K. rhizophila* or *C. glutamicum* do not function at lengths less than C_50_, suggesting that simple hydrophobicity of C_40_ or smaller carotenoids is not sufficient to mediate *K. rhizophila* or *C. glutamicum* inhibition of *C. elegans* translational surveillance. The specific color absorbance of the C_50_ carotenoids (their red color) may mediate their *C. elegans* anti-surveillance activity. For example, *C. elegans* with a translational deficit may produce blue light (Ohmiya and Hirano, 1996), and the absorption of such light by C_50_ carotenoids could suppress a light-responsive *C. elegans* surveillance pathway. Such a system might function in the dark of the soil or rotting fruit or night, but might be overwhelmed by sunlight.

Translation surveillance not only induces detoxification but also food aversion behaviors in *C. elegans*. Aversion to food associated with a toxin is an appropriate animal response since many toxins originate from bacterial pathogens that cause the rotting of food. Aversion to toxins is a common animal response since many toxins originate from a pathogen, and this response is likely to be an animal program derived from this evolutionary history. *K. rhizophila* C50 carotenoids suppressed the food aversion induced by emetogenic toxins, emetine or cisplatin. Cisplatin, which is used to block DNA replication in cancer patients, also has high emetogenic potential. Emetine, which is an antibiotic that targets eukaryotic protein synthesis, is also highly emetogenic as the name itself implies. These two drugs not only induce *C. elegans* xenobiotic detoxification but also strong food aversion behavior, both of which are strongly suppressed by bacterial C_50_ carotenoids. Chemotherapy-induced nausea and vomiting (CINV) in humans may be related to these xenobiotic aversion programs. The *K. rhizophila* C_50_ carotenoid decaprenoxanthin might for example suppress CINV, perhaps as an adjuvant in chemotherapy. We have hypothesized previously that autoimmune disorders may be triggered by reduction of function mutations in core cellular pathways such as translation that activate these surveillance and detoxification pathways (Govindan et al., 2015). Because the C_50_ carotenoid decaprenoxanthin robustly suppresses *C. elegans* translational surveillance, it may suppress the inappropriate autoimmune responses to such mutations in mammals. More broadly, bacterial gene activities that suppress animal toxin surveillance pathways could neutralize the toxicity that often plagues drug development.

## Acknowledgments

We thank members of the G.R. laboratory for helpful discussions. The work was supported by NIH Grant AG043184 (to G.R.). Some strains were provided by the CGC, which is funded by NIH Office of Research Infrastructure Programs (P40 OD010440).

---

**Supplementary Figure 1:**
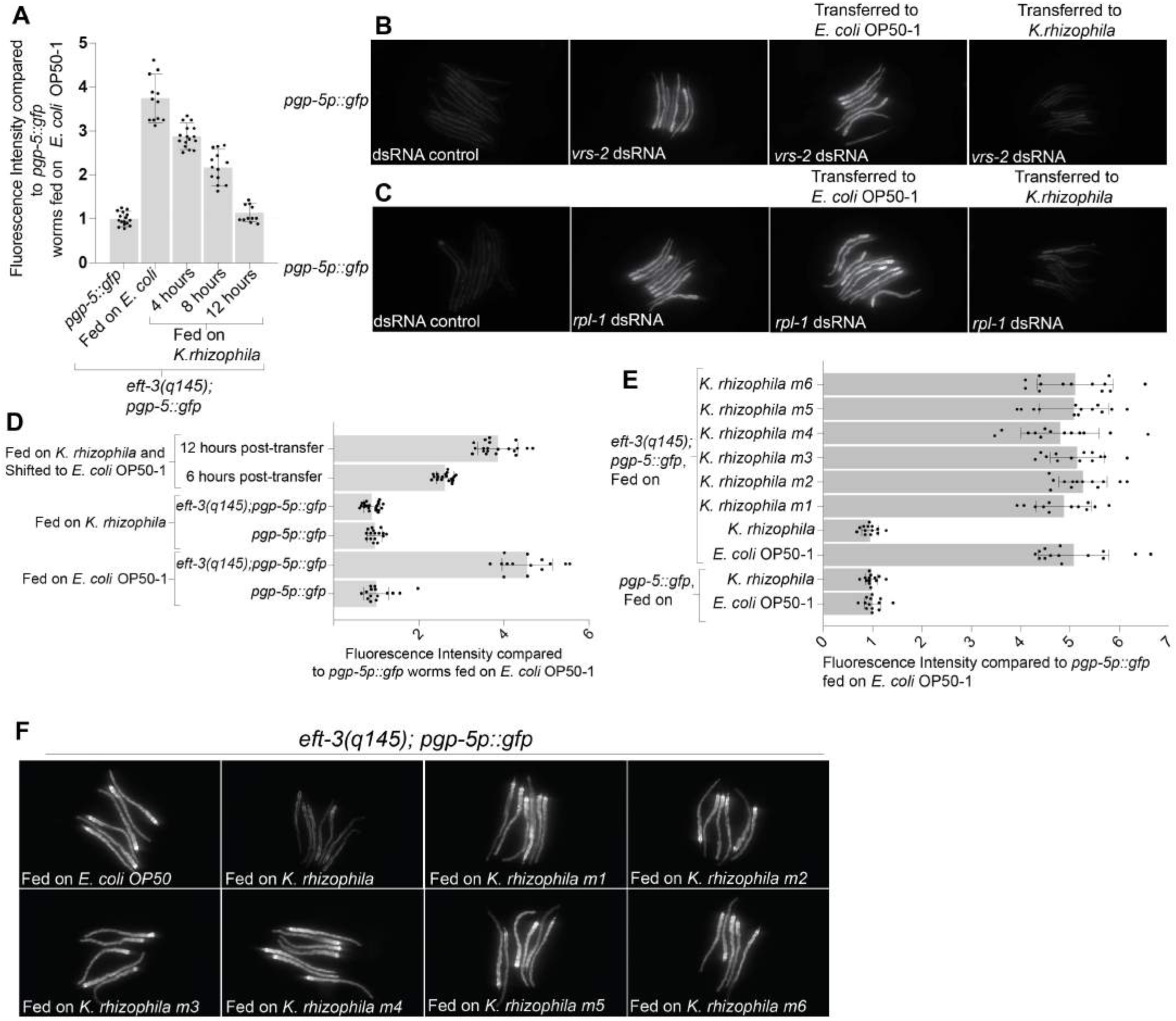
*K.rhizophila* carotenoid biosynthetic mutants fail to suppress *pgp-5::gfp* induction. A) *K. rhizophila* feeding inhibited *pgp-5p::gfp* induction in *eft-3(q145); pgp-5p::gfp* animals within 12 hours of feeding. B) While *pgp-5p::gfp* animals fed on *vrs-2* dsRNA and transferred to *E. coli* showed induction of gfp, in animals transferred to *K. rhizophila* plates, *pgp-5p::gfp* expression was reduced. C) While *pgp-5p::gfp* animals fed on *rpl-1* dsRNA and transferred to *E. coli* showed induction of gfp, in animals transferred to *K. rhizophila* plates, *pgp-5p::gfp* expression was reduced. D) *K. rhizophila* feeding suppressed *pgp-5p::gfp* induction in *eft-3(q145); pgp-5p::gfp* animals was reversible. E) *K. rhizophila* feeding reduced *pgp-5p::gfp* induction in *eft-3(q145); pgp-5p::gfp* animals significantly, while in the *K. rhizophila* mutants, *pgp-5p::gfp* expression is not affected. F) *K. rhizophila* feeding inhibited the induction of *pgp-5p::gfp* in *eft-3(q145); pgp-5p::gfp* animals while in the *K. rhizophila* mutants, *pgp-5p::gfp* expression was not affected.

**Supplementary Figure 2:**
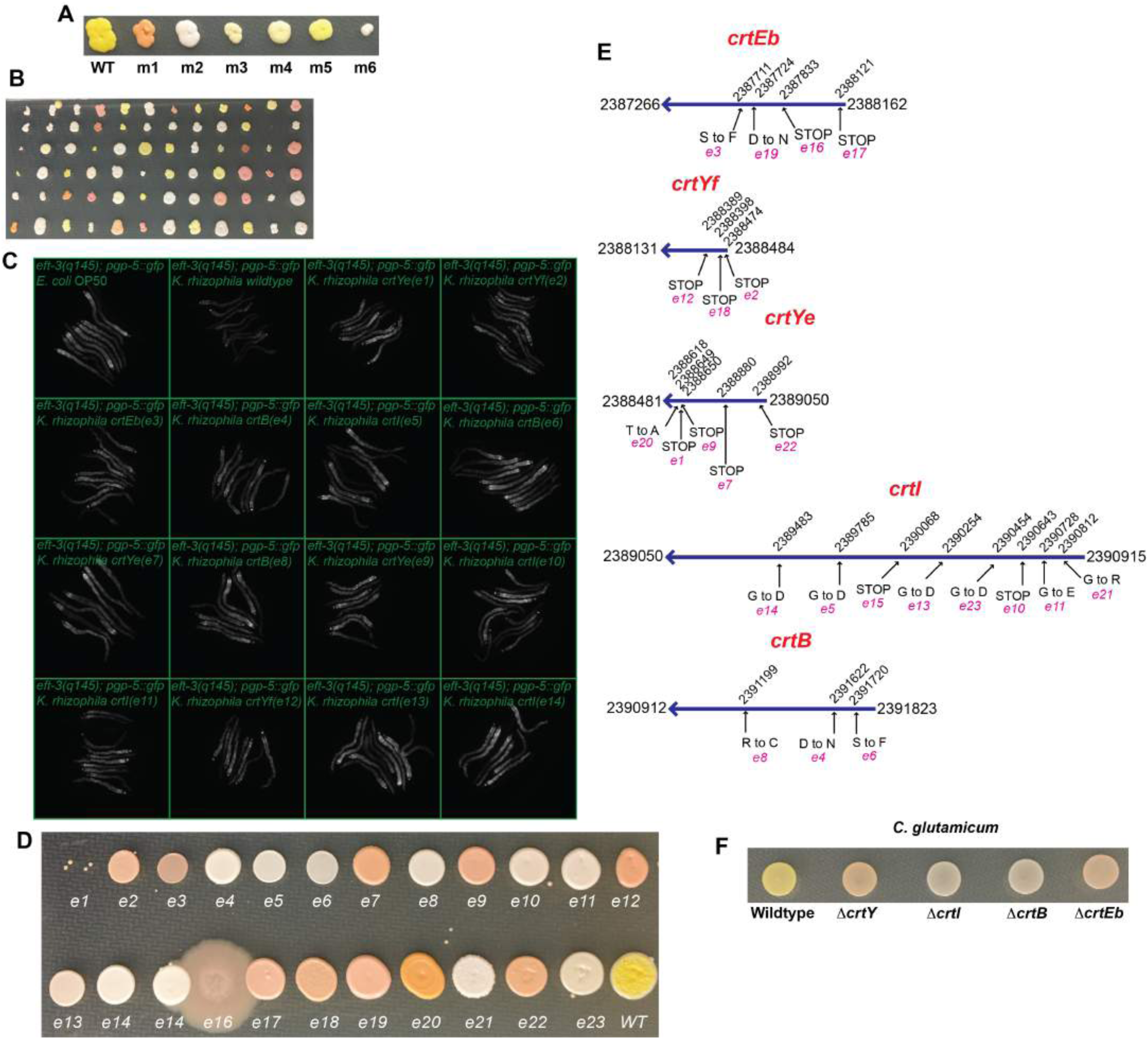
*K. rhizophila* carotenoid pathway is required for inhibition of *pgp-5p::gfp* induction in *eft-3(q145); pgp-5p::gfp* animals. A) Colony color of *K. rhizophila* wildtype was different from the colony color of the six mutants. B) Colony color of the *K. rhizophila* mutants observed from the EMS screen. C) *K. rhizophila* pigmentation mutants failed to suppress the induction of *pgp-5p::gfp* in *eft-3(q145); pgp-5p::gfp* animals. D) The pigmentation defect in 23 genome sequenced mutants. E) Predicted *K. rhizophila* carotenoid (*crt*) biosynthetic genes: *crtE* (KRH_20850; encoding GGPP synthase), *crtB* (KRH_20840; encoding phytoene synthase), *crtI* (KRH_20830; encoding phytoene desaturase), *crtEb* (KRH_20800; encoding lycopene elongase), *crtYe* (KRH_20820; encoding C_50_ carotenoid epsilon cyclase subunit) and *crtYf* (KRH_20810; encoding C_50_ carotenoid epsilon cyclase subunit). The crtYe and crtYf subunits form a heterodimeric cyclase complex that catalyzes generation of C_50_ carotenoid are shown with the genomic locus coordinates. The genomic location of the mutation with the type of change in amino acid is shown for each mutant. F) Pigmentation defects observed from *C. glutamicum* wildtype, D*crtEb*, D*crtI*, D*crtY*, and D*crtB* mutants.

**Supplementary Figure 3:**
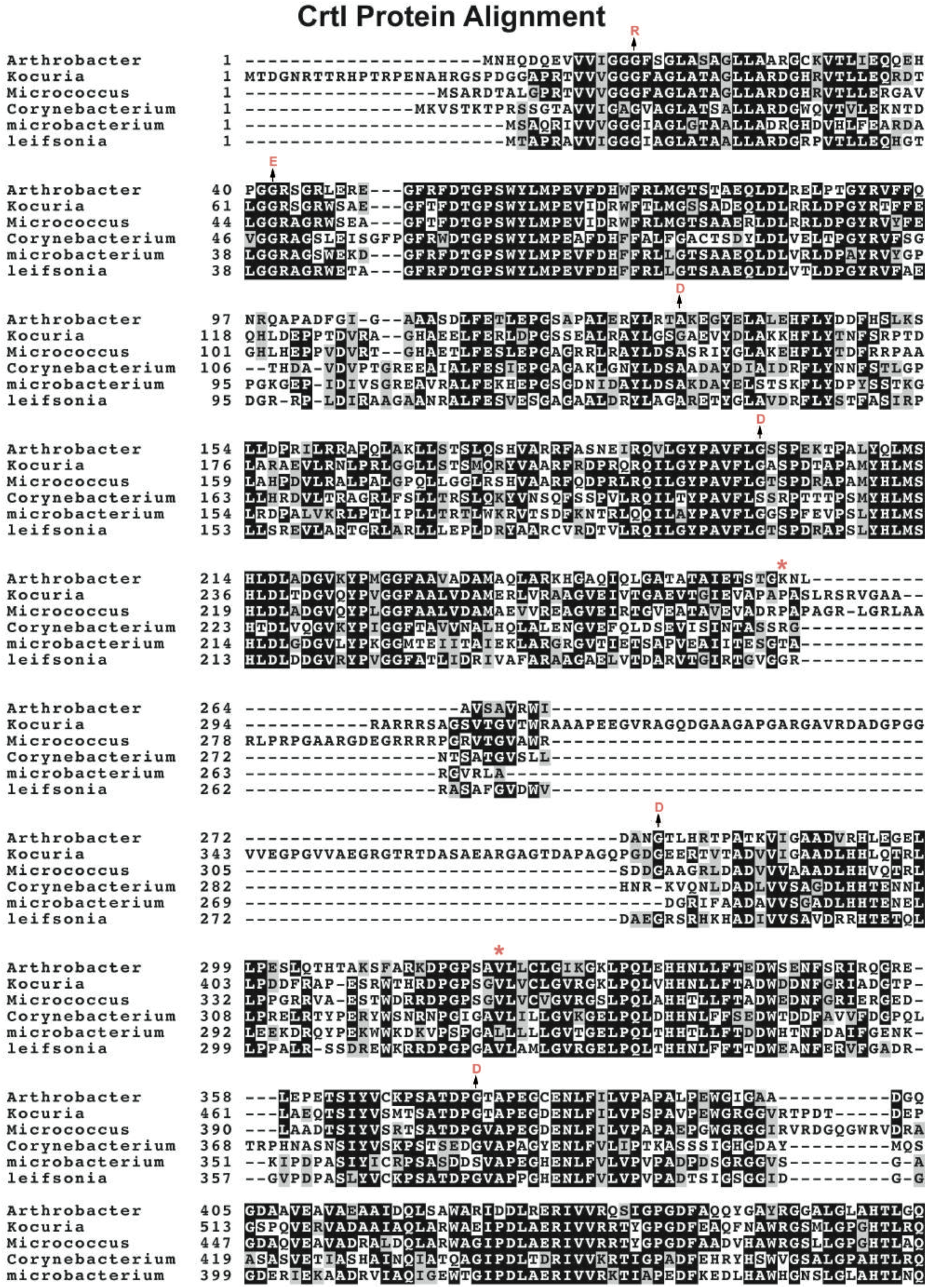
Alignment of CrtI protein in different species. Alignment of CrtI protein in different species. The amino acid conservation across different species and the mutations isolated are shown. Protein sequences were obtained from NCBI and the sequences were aligned using CLUSTAL Omega and BOXSHADE server.

**Supplementary Figure 4:**
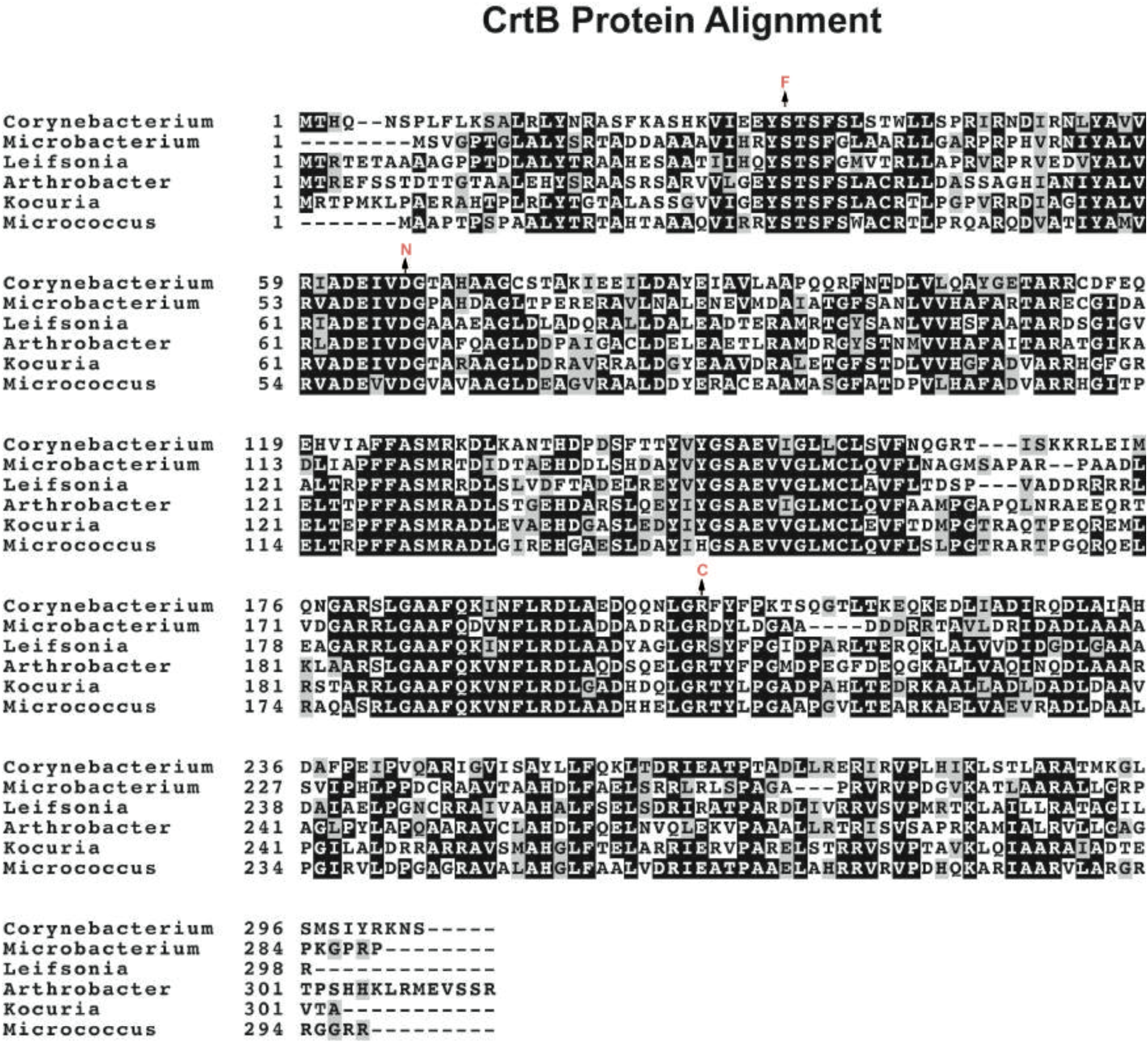
Alignment of CrtB protein in different species. Alignment of CrtB protein in different species. The amino acid conservation across different species and the mutations isolated are shown. Protein sequences were obtained from NCBI and the sequences were aligned using CLUSTAL Omega and BOXSHADE server.

**Supplementary Figure 5:**
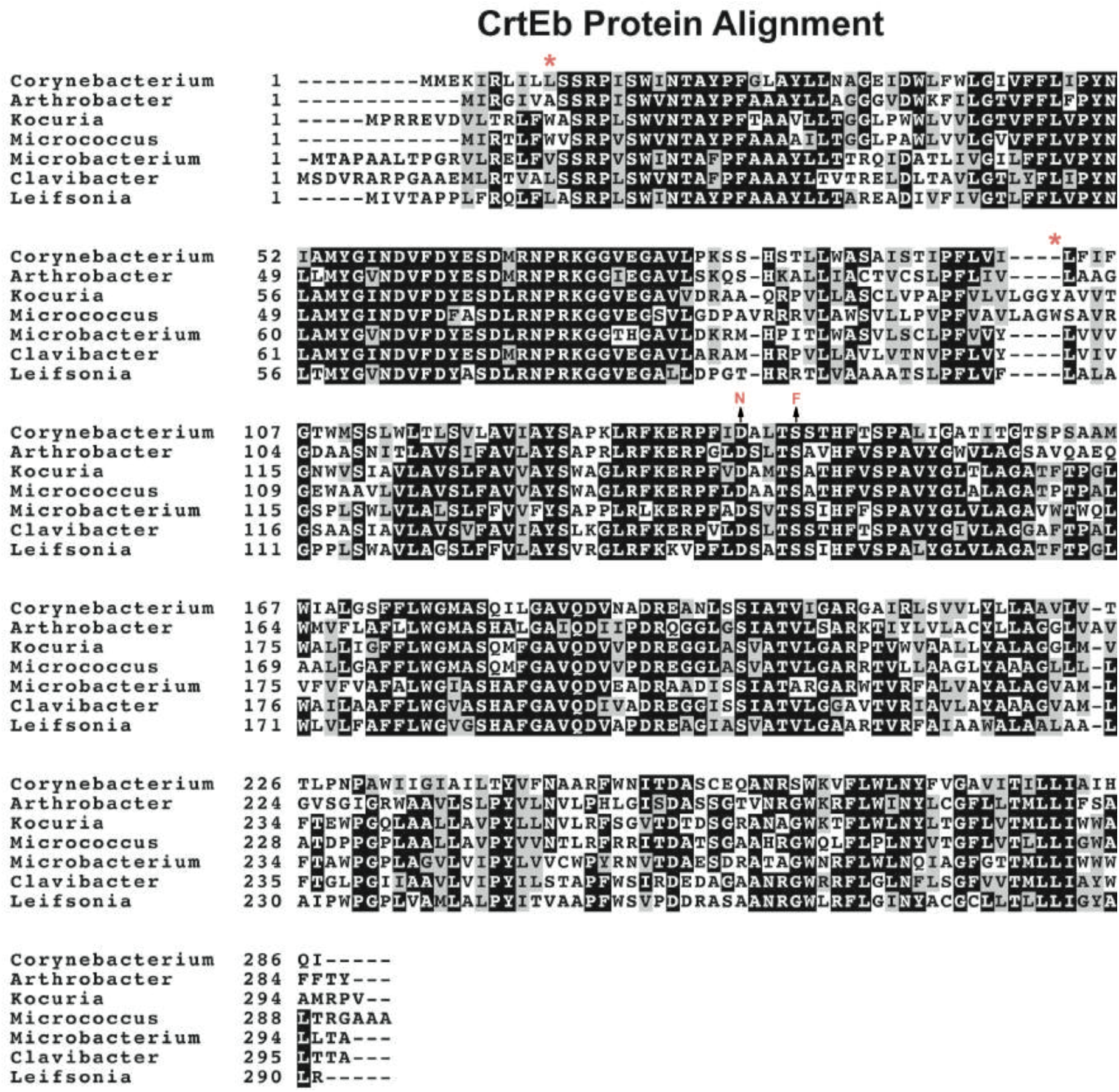
Alignment of CrtEb protein in different species. Alignment of CrtEb protein in different species. The amino acid conservation across different species and the mutations isolated are shown. Protein sequences were obtained from NCBI and the sequences were aligned using CLUSTAL Omega and BOXSHADE server.

**Supplementary Figure 6:**
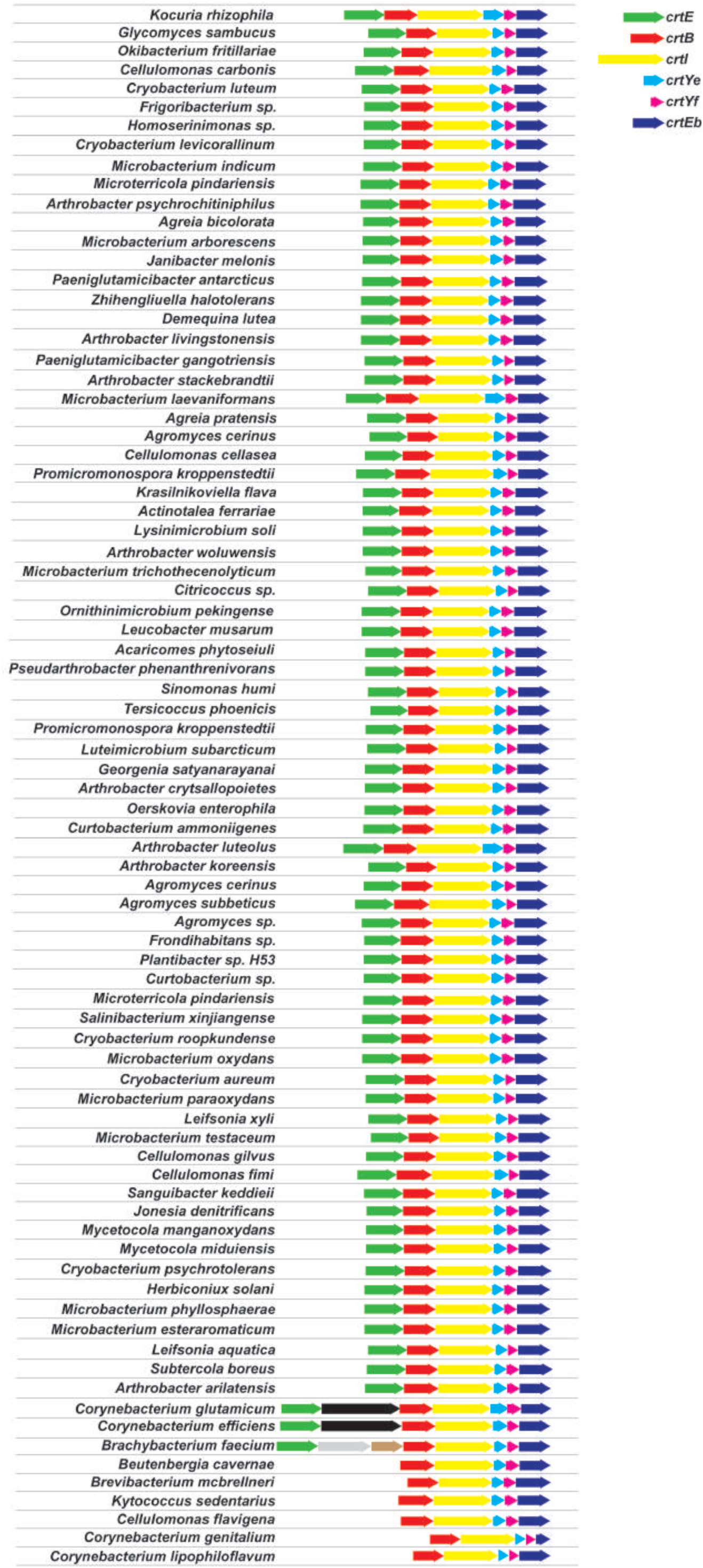
Operon structure of bacteria that contain putative gene cluster that might produce decaprenoxanthin. Genomic sequences were obtained from NCBI and analyzed for the presence of operons manually.

**Supplementary Figure 7:**
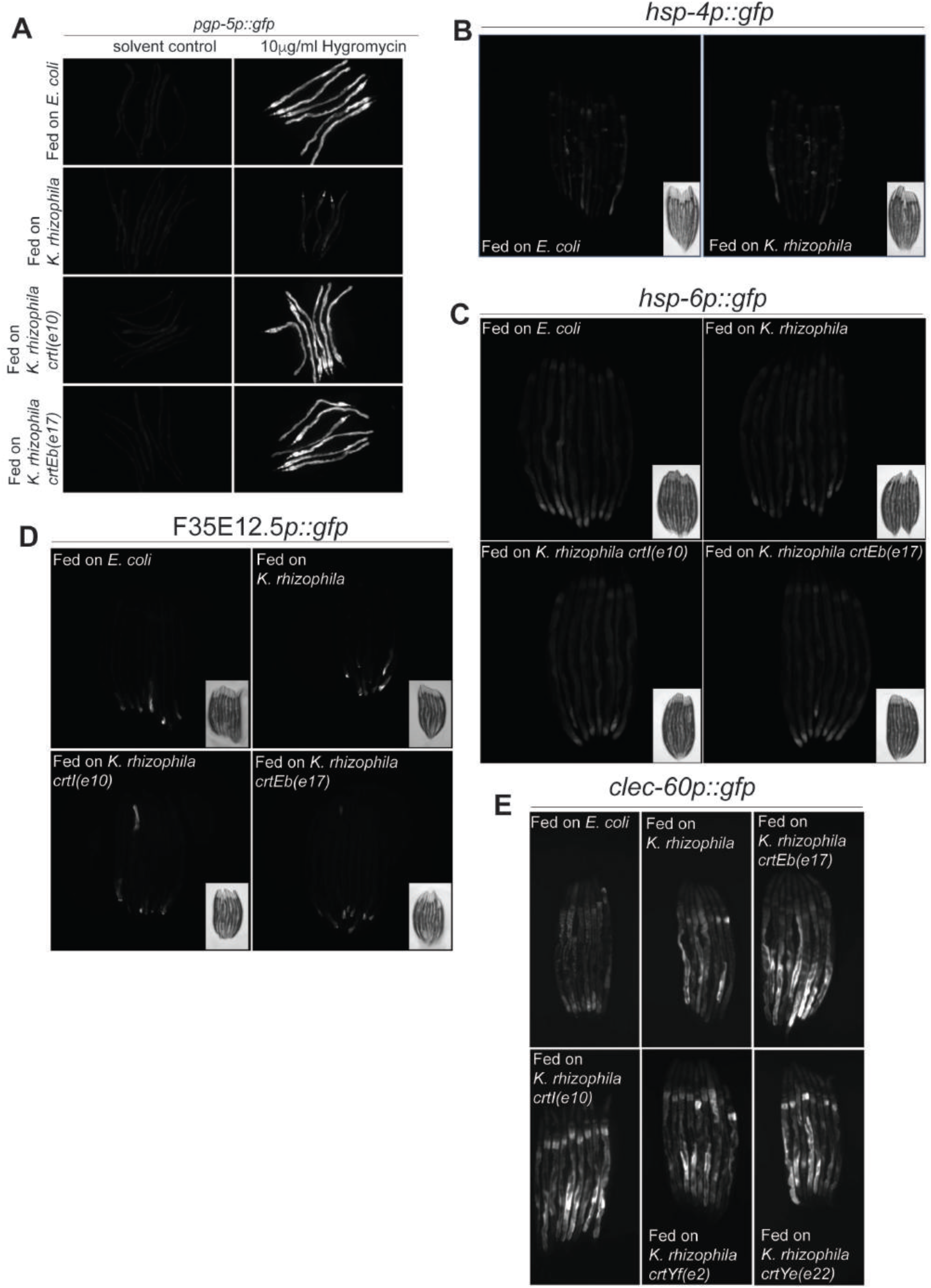
Testing whether *K. rhizophila* wildtype or carotenoid mutants regulate other stress pathways. A) *pgp-5p::gfp* induction in response to 10µg/ml of hygromycin was significantly reduced in animals grown on *K. rhizophila* wildtype while mutants in *K. rhizophila crtEb(e17)*, or *K. rhizophila crtI(e10)* did not suppress the GFP induction by hygromycin. B) *K. rhizophila* did not induce *hsp-4p::gfp*. *hsp-4p::gfp* is a reporter of mitochondrial unfolded protein response (UPR^mito^)(Calfon et al., 2002). C) *K. rhizophila* wildtype as well as *crtEb(e17)* or *crtI(e10)* mutants did not induce *hsp-6p::gfp*. *hsp-6p::gfp* is a reporter of mitochondrial unfolded protein response (UPR^mito^)(Yoneda et al., 2004). D) *K. rhizophila* wildtype as well as *crtEb(e17)* or *crtI(e10)* mutants did not induce F35E12.5p::gfp. F35E12.5p::GFP is a CUB domain protein induced by *Y. pestis*, *M. nematophilum* and *P. aeruginosa* (Bolz et al., 2010) (O’Rourke et al., 2006) (Troemel et al., 2006). E) *K. rhizophila* wildtype as well as *crtEb(e17)*, *crtI(e10)*, *crtYf(e2)* or *crtYe(e22)* mutants induced *clec-60p::gfp*. *clec-60* is a C-type lectin/CUB domain protein induced by the gram positive pathogens, *S. aureus* and *M. nematophilum* (O’Rourke et al., 2006). *clec-60::GFP* is induced by gram-positive bacteria. Because *K. rhizophila* is also a gram-positive bacterium, the induction of *clec-60* may be an immune response to a pathogen.

**Supplementary Figure 8:**
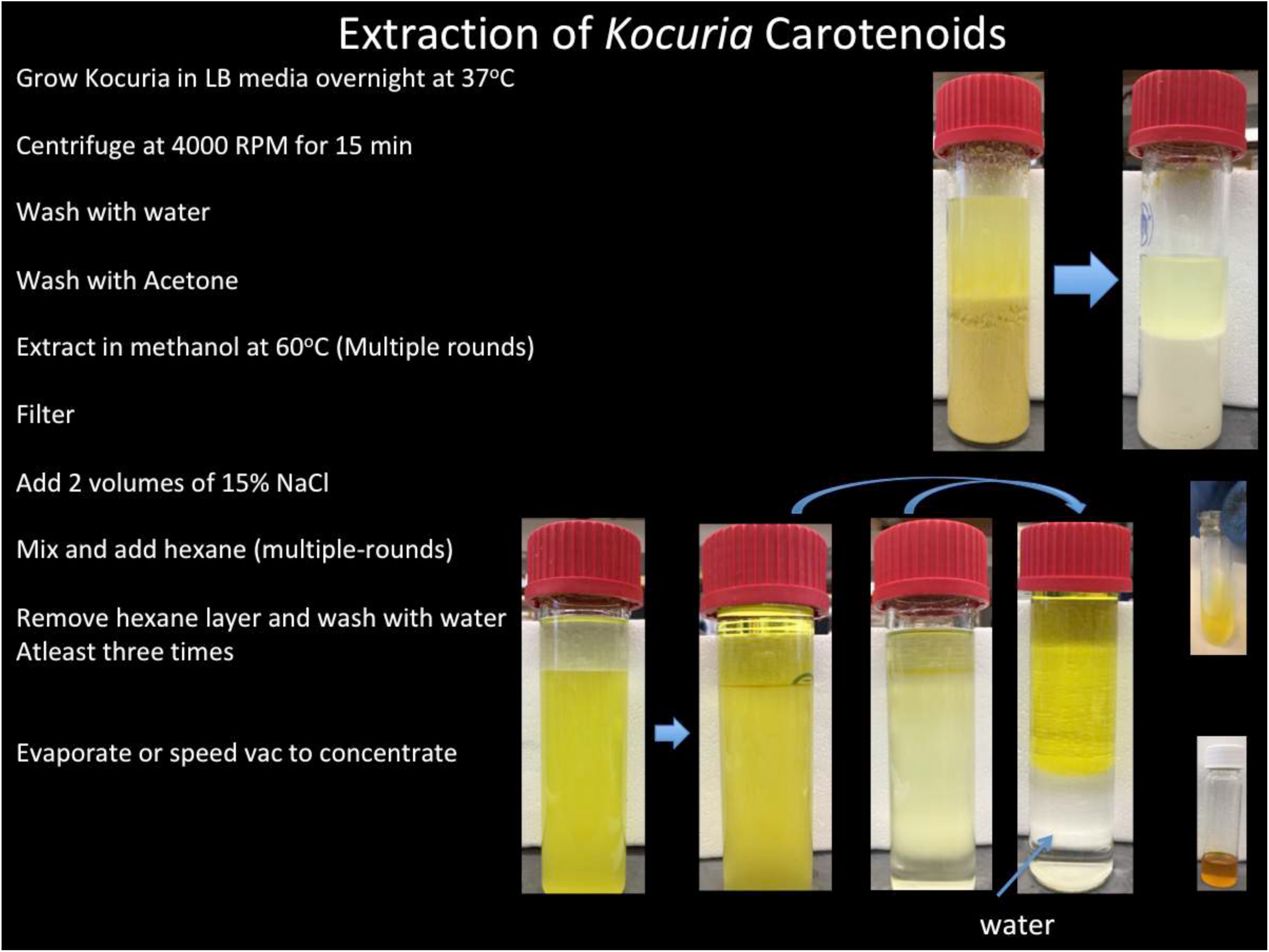
Biochemical isolation of decaprenoxanthin containing extract from *K. rhizophila*. *K. rhizophila* cultures grown in LB solution was washed with equal volume of water after centrifugation at 4000RPM for 15 min. After centrifugation to remove water, equal volume of acetone was added and centrifuged again at 4000RPM for 15 min. After removal of acetone, the bacterial pellets were extracted with methanol at 65°C in water bath after wrapping the samples with aluminum foil to protect from light. The samples were extracted with methanol multiple times. The supernatant was filtered with Whatman filter paper No1. Two-volumes of 15% sodium chloride was added to the methanol extract and after mixing equal volume of hexane was added. The yellow carotenoids were separated from the methanol-salt mix and accumulated in the hexane fraction. The hexane fraction was removed and washed at least three times with water. The hexane fraction was evaporated and the resultant carotenoid pellet was dissolved in methanol.

**Supplementary Figure 9:**
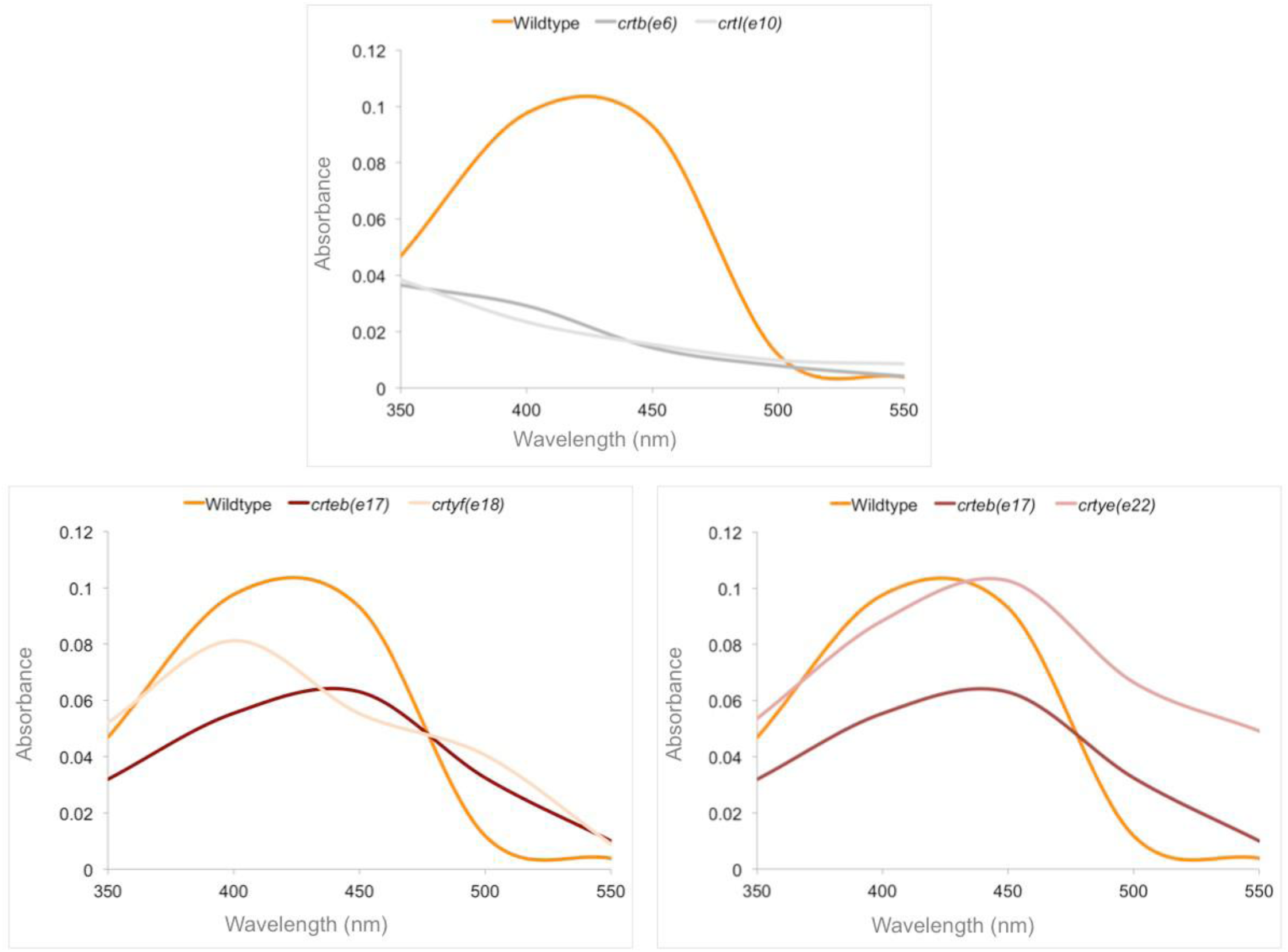
Spectrophotometric analysis of methanolic extracts. Methanolic extracts from *K. rhizophila* wildtype, *crtEb(e17)*, *crtI(e10)*, *crtb(e6)*, *crtYf(e18)*, and *crtYe(e22)* were analyzed at 350, 400, 450, 500, and 550 nm wavelengths. The absorbance of the extracts at each wavelength was plotted.

**Supplementary Figure 10:**
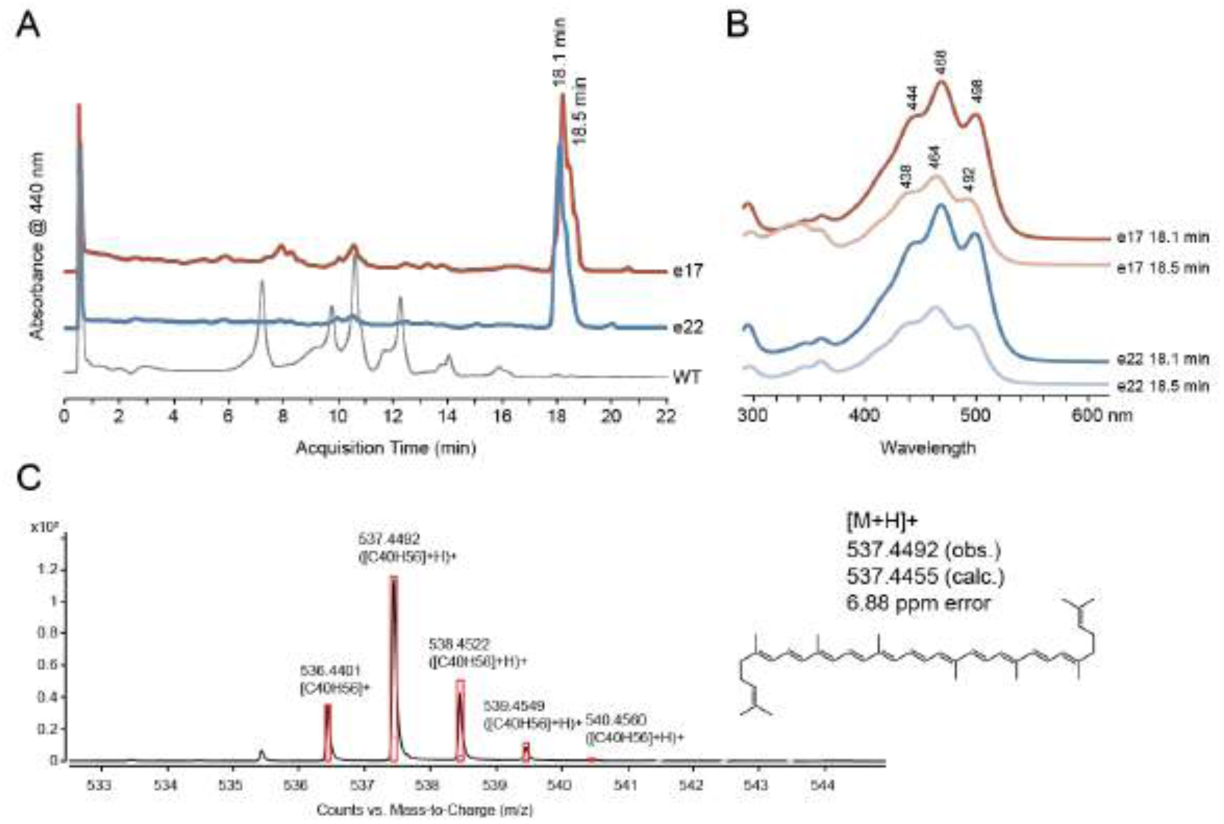
LC-MS analysis of carotenoid extracts from *K. rhizophila* mutants. A) Chromatograms of absorbance at 440 nm during reverse phase LC separation of clarified extracts on a C18 column using an elution gradient from 2% to 50% MTBE. Traces from extracts of Wildtype (gray), *crtYe(e22)* (blue), and *crtEb(e17)* (red) are overlaid. B) Absorbance spectra at the indicated elution times from *crtEb(e17)* (reds) or *crtYe(e22)* (blue) chromatograms, with absorbance peaks indicated. At least two compounds coelute in the main peak ∼18 min with distinct absorbance spectra, as shown, on the peak (18.1 min) and shoulder (18.5 min). C) MS spectral identification of a compound eluting at ∼18 min with chemical formula C40H56, consistent with lycopene, in the extracts from both mutants, as well as the calculated isotopic distribution for the identified ion (red bars).

**Supplementary Figure 11:**
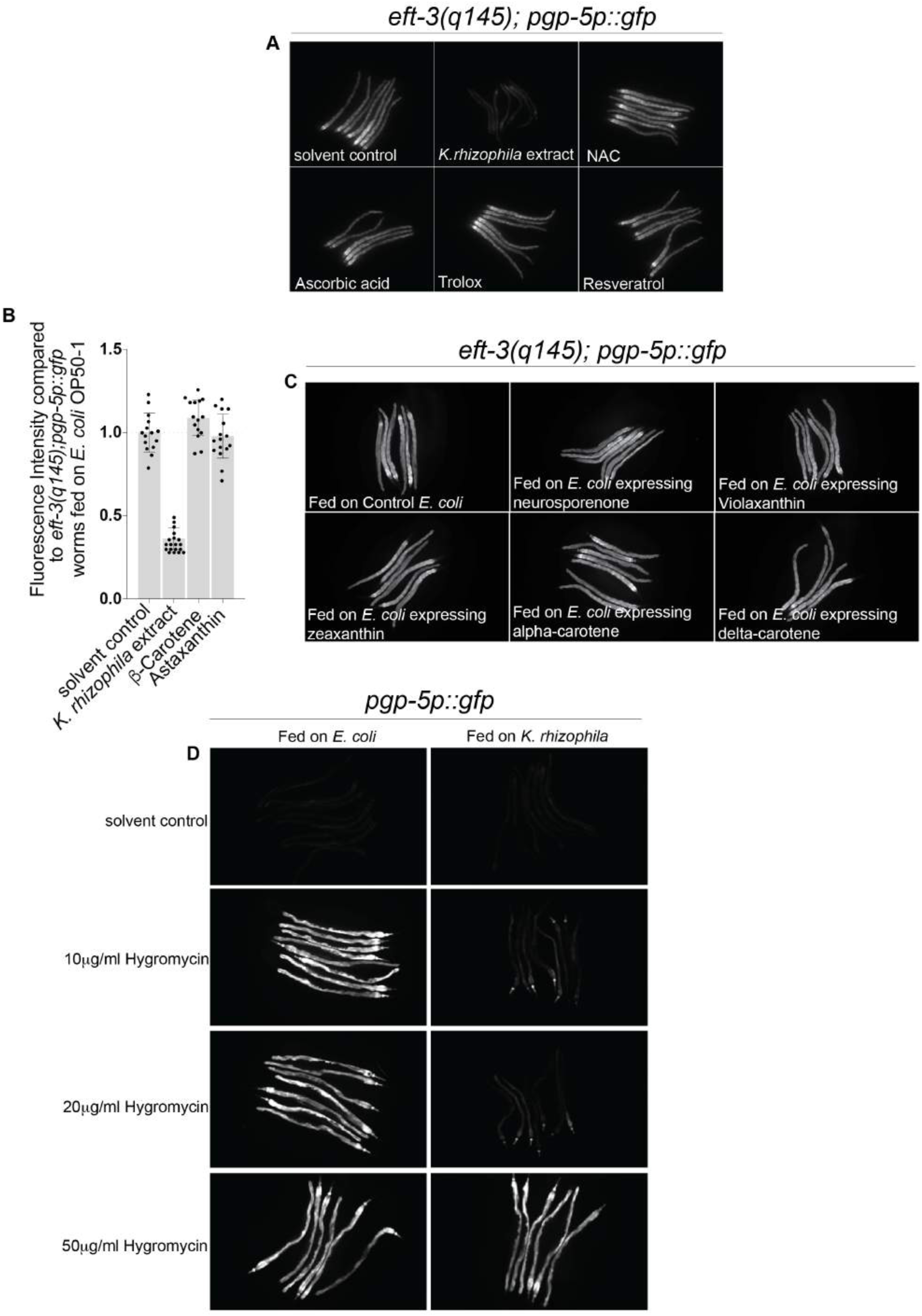
Antioxidants did not suppress *pgp-5p::gfp* induction in *eft-3(q145); pgp-5p::gfp* animals. A) N-Acetyl Cysteine, Ascorbic acid, Trolox or Resveratrol did not suppress *pgp-5p::gfp* in *eft-3(q145); pgp-5p::gfp* animals. Synchronized *eft-3(q145); pgp-5p::gfp* animals were grown from L1-larval stage in the presence of N-Acetyl Cysteine, Ascorbic acid, Trolox or Resveratrol and imaged after 50 hours at 20°C. B) Beta-carotene or Astaxanthin did not suppress *pgp-5p::gfp* in *eft-3(q145); pgp-5p::gfp* animals. Synchronized *eft-3(q145); pgp-5p::gfp* animals were grown from L1-larval stage in the presence of Beta-carotene or Astaxanthin and imaged after 50 hours at 20°C. C) *E. coli* expressing either zeaxanthin, neurosporene, violaxanthin, delta-carotene, or alpha-carotene did not suppress *pgp-5p::gfp* in *eft-3(q145); pgp-5p::gfp* animals. D) *pgp-5p::gfp* induction in response to 10µg/ml and 20µg/ml of hygromycin was significantly reduced in animals fed on *K. rhizophila* wildtype. The *pgp-5p::gfp* induction in response to 50µg/ml hygromycin is normal in animals fed on *E. coli* OP50 or on *K. rhizophila* wildtype.

**Supplementary Figure 12:**
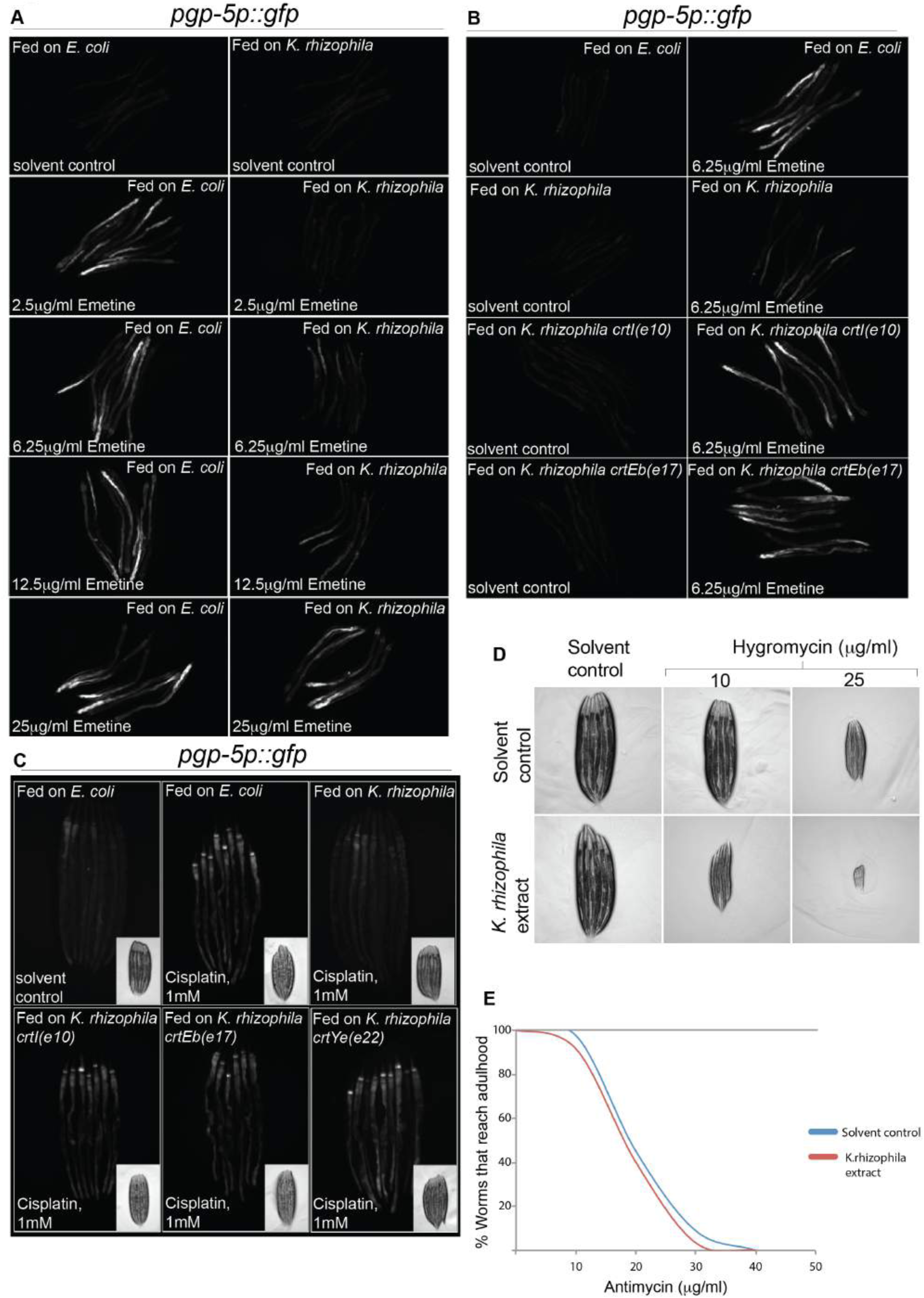
*K. rhizophila* carotenoid sensitizes the animals to toxins affecting protein synthesis but not to mitochondrial toxin. A) *pgp-5p::gfp* induction in response to 2.5µg/ml or 6.25µg/ml of emetine was significantly reduced in animals fed on *K. rhizophila* wildtype. However, in animals treated with 12.5µg/ml of emetine, *pgp-5p::gfp* induction was partially reduced in animals fed on *K. rhizophila*. In animals treated with 25µg/ml of emetine, the induction of *pgp-5p::gfp* was normal both animals treated with either *E. coli* OP50 or *K. rhizophila* wildtype. B) *pgp-5p::gfp* induction in response to 6.25µg/ml of emetine was significantly reduced in animals fed on *K. rhizophila* wildtype while mutants in *K. rhizophila crtEb(e17)* or *K. rhizophila crtI(e10)* did not suppress the GFP induction C) Animals treated with *K. rhizophila* carotenoid extract were hypersensitive to Hygromycin. Synchronized wildtype animals treated with solvent control or *K. rhizophila* extract from L1-larval stage were exposed to hygromycin at L3-larval stage and scored for development 60 hours later at 20°C. Animals treated with *K. rhizophila* extract and hygromycin appear smaller compared to animals on solvent control and hygromycin. D) *pgp-5p::gfp* induction in response to 1mM of cisplatin was significantly reduced in animals fed on *K. rhizophila* wildtype while mutants in *K. rhizophila crtEb(e17)*, *crtYe(e22)* or *crtI(e10)* did not suppress the GFP induction. Synchronized *pgp-5p::gfp* animals were grown on *E. coli* OP50 plates from L1-larval stage and transferred to either *K. rhizophila* wildtype or *K. rhizophila crtEb(e17)*, *crtYe(e22)* or *crtI(e10)* at L4-stage at 20°C. After 24 hours, the animals were treated with either solvent or cisplatin and scored for induction of GFP after 24 hours. E) Animals fed on control extract or *K. rhizophila* extract were sensitive to antimycin. Synchronized wildtype animals treated with solvent control or *K. rhizophila* extract from L1-larval stage were exposed to antimycin at L3-larval stage and scored for development 60 hours later at 20°C.

**Supplementary Table 1.** Analysis of the effects of *K. rhizophila* mutants on *gfp* induction in *eft-3(q145); pgp-5p::gfp* animals and *pgp-5p::gfp* animals.

| ***Kocuria***<br>**genotype** | **% *eft-3(q145); pgp-5::gfp* animals showing**<br>**GFP induction** |  | **% *pgp-5::gfp* animals**<br>**showing GFP**<br>**induction** |  |
| --- | --- | --- | --- | --- |
|  | **%** | **N** | **%** | **N** |
| Wildtype | 0 | 26 | 0 | 24 |
| *crtYe(e1)* | 95.6 | 23 | 0 | 24 |
| *crtYf(e2)* | 94.4 | 36 | 0 | 23 |
| *crtEb(e3)* | 76.6 | 30 | 0 | 24 |
| *crtB(e4)* | 80.6 | 31 | 0 | 24 |
| *crtl(e5)* | 95.8 | 24 | 0 | 27 |
| *crtB(e6)* | 92.5 | 27 | 0 | 29 |
| *crtYe(e7)* | 93.3 | 45 | 0 | 24 |
| *crtB(e8)* | 95.8 | 24 | 0 | 23 |
| *crtYe(e9)* | 95.6 | 23 | 0 | 24 |
| *crtl(e10)* | 100 | 24 | 0 | 24 |
| *crtl(e11)* | 95.8 | 24 | 0 | 27 |
| *crtYf(e12)* | 100 | 27 | 0 | 29 |
| *crtl(e13)* | 100 | 29 | 0 | 31 |
| *crtl(e14)* | 100 | 31 | 0 | 23 |
| *crtl(e15)* | 95.6 | 23 | 0 | 36 |
| *crtEb(e16)* | 94.4 | 36 | 0 | 30 |
| *crtEb(e17)* | 76.6 | 30 | 0 | 31 |
| *crtYf(e18)* | 80.6 | 31 | 0 | 24 |
| *crtEb(e19)* | 95.8 | 24 | 0 | 27 |
| *crtYe(e20)* | 92.5 | 27 | 0 | 45 |
| *crtl(e21)* | 93.3 | 45 | 0 | 24 |
| *crtYe(e22)* | 95.8 | 24 | 0 | 23 |
| *crtl(e23)* | 95.6 | 23 | 0 | 24 |
| *m24* | 100 | 24 | 0 | 24 |
| *m25* | 95.8 | 24 | 0 | 27 |
| *m26* | 100 | 27 | 0 | 29 |
| *m27* | 100 | 29 | 0 | 24 |
| *m28* | 100 | 24 | 0 | 27 |
| *m29* | 96.2 | 27 | 0 | 23 |
| *m30* | 95.6 | 23 | 0 | 24 |
| *m31* | 95.6 | 23 | 0 | 24 |
| *m32* | 94.4 | 36 | 0 | 27 |
| *m33* | 76.6 | 30 | 0 | 29 |
| *m34* | 80.6 | 31 | 0 | 24 |
| *m35* | 95.8 | 24 | 0 | 27 |
| *m36* | 92.5 | 27 | 0 | 23 |
| *m37* | 93.3 | 45 | 0 | 23 |
| *m38* | 95.8 | 24 | 0 | 24 |
| *m39* | 95.6 | 23 | 0 | 24 |
| *m40* | 100 | 24 | 0 | 27 |
| *m41* | 95.8 | 24 | 0 | 29 |
| *m42* | 100 | 27 | 0 | 24 |
| *m43* | 100 | 29 | 0 | 27 |
| *m44* | 95.6 | 23 | 0 | 23 |
| *m45* | 95.6 | 23 | 0 | 27 |
| *m46* | 94.4 | 36 | 0 | 24 |
| *m47* | 76.6 | 30 | 0 | 23 |
| *m48* | 80.6 | 31 | 0 | 24 |
| *m49* | 95.8 | 24 | 0 | 24 |
| *m50* | 92.5 | 27 | 0 | 27 |
| *m51* | 93.3 | 45 | 0 | 29 |
| *m52* | 95.8 | 24 | 0 | 31 |
| *m53* | 95.6 | 23 | 0 | 23 |
| *m54* | 100 | 24 | 0 | 36 |
| *m55* | 95.8 | 24 | 0 | 24 |
| *m56* | 100 | 27 | 0 | 23 |
| *m57* | 100 | 29 | 0 | 24 |
| *m58* | 95.8 | 24 | 0 | 24 |
| *m59* | 95.6 | 23 | 0 | 27 |
| *m60* | 94.4 | 36 | 0 | 29 |
| *m61* | 76.6 | 30 | 0 | 31 |
| *m62* | 80.6 | 31 | 0 | 23 |
| *m63* | 95.8 | 24 | 0 | 36 |
| *m64* | 92.5 | 27 | 0 | 30 |
| *m65* | 93.3 | 45 | 0 | 31 |
| *m66* | 95.8 | 24 | 0 | 24 |
| *m67* | 95.6 | 23 | 0 | 27 |
| *m68* | 100 | 24 | 0 | 45 |
| *m69* | 100 | 29 | 0 | 24 |
| *m70* | 95.8 | 24 | 0 | 23 |
| *m71* | 95.6 | 23 | 0 | 24 |
| *m72* | 0 | 29 | 0 | 24 |
| *m73* | 0 | 23 | 0 | 27 |
| *m74* | 4.3 | 23 | 0 | 29 |
| *m75* | 3.3 | 30 | 0 | 24 |
| *m76* | 3.2 | 31 | 0 | 24 |
| *m77* | 0 | 24 | 0 | 23 |
| *m78* | 0 | 27 | 0 | 24 |
| *m79* | 2.2 | 45 | 0 | 24 |
| *m80* | 0 | 24 | 0 | 27 |
| *m81* | 0 | 30 | 0 | 29 |
| *m82* | 0 | 31 | 0 | 31 |
| *m83* | 0 | 24 | 0 | 23 |
| *m84* | 0 | 27 | 0 | 36 |
| *m85* | 0 | 45 | 0 | 30 |
| *m86* | 0 | 23 | 0 | 31 |
| *m87* | 0 | 23 | 0 | 24 |
| *m88* | 3.3 | 30 | 0 | 27 |
| *m89* | 0 | 31 | 0 | 45 |
| *m90* | 0 | 24 | 0 | 24 |
| *m91* | 0 | 27 | 0 | 24 |
| *m92* | 0 | 45 | 0 | 23 |
| *m93* | 0 | 24 | 0 | 24 |
| *m94* | 0 | 30 | 0 | 24 |
| *m95* | 0 | 29 | 0 | 27 |
| *m96* | 3.7 | 27 | 0 | 29 |

## Materials and methods

N2 Bristol was the wildtype strain used.

The following strains and mutant alleles were used:

SJ4100 [*zcIs13[hsp-6::GFP*],

WE5172 [*ajIs1(pgp-5::gfp)X*],

JG16 [*eft-3(q145)/hT2[bli-4(e937) let-?(q782) qIs48] (I;III); ajIs1(pgp-5::gfp)X*],

JG20 [*ajIs1(pgp-5::gfp)*; *agEX(pvha-6::mcherry::zip-2+ myo-2::gfp)*],

AY101 [*acls101[F35E12.5P::GFP + rol-6(su1006)*]

SJ4005 [*zcIs4 [hsp-4::GFP] V*],

SJ4100(*zcIs13[hsp-6::GFP]*)

### Growth and handling of microbes used

16S ribosomal sequence was amplified using specific primers and sequenced to identify the microbes. LB media as well as plates was used for culturing *Kocuria rhizophila*, *Arthrobacter arilaitensis* and *Corynebacterium glutamicum a*nd its mutants. 500 µl of overnight culture was seeded onto SK media plates and incubated at room temperature for 2 days before initiating the experiments. For experiments involving *Kocuria rhizophila* wildtype and mutants, *Arthrobacter arilaitensis*, and *Corynebacterium glutamicum* wildtype and as well as mutants, synchronized L1-larval stage animals grown until L4-larval stage or day one of adulthood in *E. coli* OP50 seeded plates and was washed in M9 buffer at least five times before transferring to the appropriate bacterial food.

### Drug treatments

Hygromycin diluted in M9 solution to the desired concentration was added onto preseeded *E.coli* OP50 bacteria containing NGM plates. Stock solution of emetine or cisplatin was diluted in M9 and the desired concentration was added onto preseeded *E.coli* OP50 bacteria containing NGM plates. 750µg/ml of *K. rhizophila* extracts was added onto preseeded *E.coli* OP50 bacteria containing NGM plates containing appropriate concentrations of hygromycin or cisplatin or emetine. For the xenobiotic experiments, synchronized L1-stage animals were dropped onto the drug containing plates and scored 4 days later.

Food aversion assay was performed by adding either hygromycin or cisplatin to *E. coli* OP50 lawns that contained synchronized L4 larval stage worms that were treated with 750µg/ml of *K. rhizophila* extracts. Aversion index was calculated by counting number of animals off lawn over number of total animls counted at desired time point (Aversion index=N_(animals off lawn)_/N_(total animals)_. Each experiment was conducted in triplicate and the experiments were repeated in three independent trials.

### RNAi Assays

For RNAi assays synchronized L1 larval stage animals of the appropriate genotype were fed with appropriate RNAi clones until they reach day one of adulthood. Subsequently, the RNAi-treated animals were washed in M9 at least five times to remove the *E. coli* bacteria and transferred to *K. rhizophila* seeded plates or *E. coli* OP50 seeded plates.

### Microscopy

Nematodes were mounted onto agar pads and images were acquired using a Zeiss AXIO Imager Z1 microscope fitted with a Zeiss AxioCam HRm camera and Axiovision 4.6 (Zeiss) software. All the fluorescence images shown within the same figure panel were collected together using the same exposure time. Images were converted to 8-bit image, thresholded and quantified using ImageJ. Student’s t test was used determine statistical significance. Low-magnification bright-field and GFP fluorescence images were acquired using a Zeiss AxioZoom V16, equipped with a Hammamatsu Orca flash 4.0 digital camera, and using Axiovision ZEN software.

### Multiple alignment of protein sequences

Multiple alignments were performed using Clustal Omega: (https://www.ebi.ac.uk/Tools/msa/clustalo/)

### *K. rhizophila* EMS mutagenesis screen

Mutagenesis was performed by treatment of overnight culture of *K. rhizophila* in PBS solution with 50 mM EMS for 45 minutes at 37°C. Serial dilutions of the mutagenized *K. rhizophila* cultures were plated onto LB media plates and ∼2000 mutagenized bacterial colonies were picked and grown in LB solution. 500 µl of overnight culture was seeded onto SK media plates and incubated at room temperature for two days before initiating the experiments. Synchronized L1-larval stage in *eft-3(q145);pgp-5p::gfp* animals grown until L4-larval stage or day one of adulthood in *E. coli* OP50 seeded plates and was washed in M9 buffer at least five times before transferring to the *K. rhizophila* mutant bacterial food. The plates were visually screened after 24 hours for GFP induction.

### Identification of EMS induced mutations by whole genome sequencing

Genomic DNA extraction, library prep, Illumina MiniSeq sequencing and bioinformatics were all performed by The Sequencing Center (www.thesequencingcenter.com). For genomic DNA isolation, The Sequencing Center used the Quick-DNA Fungal/Bacterial Miniprep Kit, a standard DNA extraction kit that uses bead beating methods to dissolve cell walls without organic denaturants or proteinases. The Illumina Nextera XT DNA Library Prep kit and protocol were used for library preparation. DNA quantitation was performed on a Qubit Fluorometer. The Illumina MiniSeq System Denature and Dilute Libraries Guide was followed to dilute and denature the final library pool, including a 1% PhiX spike-in control before loading onto the sequencer. The final library was loaded onto an Illumina High Output Reagent Kit on an Illumina MiniSeq with 2 x 150 paired-end reads. Sequencing data was processed through a Geneious Prime bioinformatics pipeline. Sequencing reads were aligned to the *Kocuria rhizophila* DC2201 (NC_010617.1) reference genome to identify mutant alleles and gene variants.

### Isolation of *K. rhizophila* carotenoids

Carotenoid isolation from *K. rhizophila* was isolated as described^1^ with the following modifications. *K. rhizophila* cultures grown in LB solution was washed with equal volume of water after centrifugation at 4000RPM for 15 min. After centrifugation to remove water, equal volume of acetone was added and centrifuged again at 4000RPM for 15 min. After removal of acetone, the bacterial pellets were extracted with methanol at 65°C in water bath after wrapping the samples with aluminum foil to protect from light. The samples were extracted with methanol multiple times. The supernatant was filtered with Whatman filter paper No1. Two-volumes of 15% sodium chloride was added to the methanol extract and after mixing equal volume of hexane was added. The yellow carotenoids were separated from the methanol-salt mix and accumulated in the hexane fraction. The hexane fraction was removed and washed at least three times with water. The hexane fraction was evaporated and the resultant carotenoid pellet was dissolved in methanol.

### High performance liquid chromatography

Crude methanolic extracts were separated over an Agilent Eclipse Plus C18 4.6 x 250 mm column with a 5-micron particle size using an Agilent 1200 HPLC equipped with a diode array detector, autosampler, column oven, solvent degasser, and binary pump. The mobile phases were (A) water vs. (B) methanol at a flow rate of 2 mL/min. The column was preequilibrated at 40°C with 90% B prior to sample injection. Following injection, the column was washed isocratically for 5 min at 90% B before ramping to 100% B over 5 min. Eluate absorbance spectra were monitored from 300-700 nm.

### Liquid chromatography mass spectrometry (LC-MS)

Methanol (MeOH) extracts were diluted in dichloromethane (DCM) and filtered over prewashed silica gel using 15:85 methanol/DCM. The visibly colored eluate was collected, aliquoted, and dried under vacuum. Dry aliquots were stored under Ar at −30 °C and were resuspended just prior to analysis in a minimal volume of 1:9 water/MeOH. The material was separated over Waters XBridge C18 1 x 100 mm column with a 3.5-micron particle size at 25 °C using an Agilent 1200 HPLC equipped with a diode array detector, autosampler, column oven, solvent degasser, and binary pump. The mobile phases were (A) LC-MS grade 1:9 water/MeOH vs. (B) HPLC-grade methyl tert-butyl ether (MTBE) at a flow rate of 150 µL/min. The column was preequilibrated with 2% B prior to sample injection. Following injection, the column was washed isocratically for 5 min at 2% B before ramping to 30% B over 25 min. Eluate absorbance spectra were monitored from 225-650 nm, and high-resolution mass analysis was performed on-line with an Agilent 6230 TOF mass spectrometer equipped with a Multimode source in positive atmospheric pressure chemical ionization (APCI+) mode. The MS source parameters were as follows: nitrogen drying gas temperature, 325 °C; drying gas flow, 5 L/min; vaporizer temperature, 200 °C; charging voltage, 2000 V; capillary voltage, 2000 V; nebulizer pressure, 25 psig; and corona current, 4 µA. The fragmentor voltage was set to 250 V, and MS1 scans were acquired from 100 – 3200 m/z at a rate of 1 Hz. Compound peaks were identified by searching high resolution mass spectra by chemical formula from a database of known carotenoids, using a stringent mass error threshold (5 ppm), and then correlating extracted ion chromatograms (EIC) of candidate hits to 440 nm absorbance elution peak profiles.

### Putative “decaprenoxanthin” carotenoid biosynthetic cluster from microbes

Putative carotenoid biosynthetic cluster of *Leifsonia xyli* (Lxx15630, Lxx15620, Lxx15610, Lxx15600, Lxx15590, and Lxx15580)^3^, *Microbacterium testaceum* (MTES_3133, MTES_3132, MTES_3131, MTES_3130, MTES_3129, and MTES_3128)^4^, *Cellvibrio gilvus* (Celgi_1516, Celgi_1515, Celgi_1514, Celgi_1513, Celgi_1512, and Celgi_1511)^5^, *Cellulomonas fimi* (Celf_3171, Celf_3170, Celf_3169, Celf_3168, Celf_3167, and Celf_3166)^5^, *Sanguibacter keddieii* (Sked_12750, Sked_12760, Sked_12770, Sked_12780, Sked_12790, and Sked_12800)^6^, *Jonesia denitrificans* (Jden_0342, Jden_0341, Jden_0340, Jden_0339, Jden_0338, and Jden_0337)^7^, *Mycetocola manganoxydans* (D9V29_RS08865, D9V29_RS08870, D9V29_RS08875, D9V29_RS08880, D9V29_RS08885, and D9V29_RS08890), *Mycetocola miduiensis* (BM197_RS02470, BM197_RS02475, BM197_RS02480, BM197_RS02485, BM197_RS02490, and BM197_RS02495), *Cryobacterium psychrotolerans* (BLQ39_RS02180, BLQ39_RS02185, BLQ39_RS02190, BLQ39_RS02195, BLQ39_RS02200, and BLQ39_RS02205), *Subtercola boreus* (B7R21_RS02695, B7R21_RS02700, B7R21_RS02705, B7R21_RS02710, B7R21_RS02715, and B7R21_RS02720), *Herbiconiux solani* (HSO01S_RS07000, HSO01S_RS07005, HSO01S_RS07010, HSO01S_RS07015, HSO01S_RS07020, and HSO01S_RS07025), *Microbacterium phyllosphaerae* (D3H67_RS09120, D3H67_RS09125, D3H67_RS09130, D3H67_RS09135, D3H67_RS09140, and D3H67_RS09145), *Leifsonia aquatica* (N136_RS22055, N136_RS22060, N136_RS22065, N136_RS22070, N136_RS22075, and N136_RS22080), *Microbacterium esteraromaticum* (B4U78_RS09520, B4U78_RS09525, B4U78_RS09530, B4U78_RS09535, B4U78_RS09540, and B4U78_RS09545), *Plantibacter sp.* H53 (A4X17_RS18565, A4X17_RS18570, A4X17_RS18575, A4X17_RS18580, A4X17_RS18585, and A4X17_RS18590), *Curtobacterium sp*. (ASF23_RS14315, ASF23_RS14320, ASF23_RS14325, ASF23_RS14330, ASF23_RS14335, and ASF23_RS14340), *Microterricola pindariensis* (GY24_RS04745, GY24_RS04750, GY24_RS04755, GY24_RS04760, GY24_RS04765, and GY24_RS04770), *Frondihabitans sp.* (EDF46_RS08000, EDF46_RS08005, EDF46_RS08010, EDF46_RS08015, EDF46_RS08020, and EDF46_RS08025), *Salinibacterium xinjiangense* (SAMN06296378_0676, SAMN06296378_0677, SAMN06296378_0678, SAMN06296378_0679, SAMN06296378_0680, and SAMN06296378_0681), *Agromyces sp.* (AVP42_RS01110, AVP42_RS01115, AVP42_RS01120, AVP42_RS01125, AVP42_RS01130, and AVP42_RS01135), *Microbacterium barkeri* (MBR4_RS00310, MBR4_RS00315, MBR4_RS00320, MBR4_RS00325, MBR4_RS00330, and MBR4_RS00335), *Arthrobacter koreensis* (BN2404_RS04370, BN2404_RS04375, BN2404_RS04380, BN2404_RS04385, BN2404_RS04390, and BN2404_RS04395), *Cryobacterium roopkundense* (GY21_RS00565, GY21_RS00570, GY21_RS00575, GY21_RS00580, GY21_RS00585, and GY21_RS00590), *Microbacterium oxydans* (RN51_RS07325, RN51_RS07330, RN51_RS07335, RN51_RS07340, RN51_RS07345, and RN51_RS07350), *Arthrobacter luteolus* (AL3_RS02570, AL3_RS02575, AL3_RS02580, AL3_RS02585, AL3_RS02590, and AL3_RS02595), *Cryobacterium aureum* (CJ028_RS03575, CJ028_RS03580, CJ028_RS03585, CJ028_RS03590, CJ028_RS03595, and CJ028_RS03600), *Curtobacterium ammoniigenes* (CAM01S_RS13455, CAM01S_RS13460, CAM01S_RS13465, CAM01S_RS13470, CAM01S_RS13475, and CAM01S_RS13480), *Oerskovia enterophila* (OJAG_RS08920, OJAG_RS08925, OJAG_RS08930, OJAG_RS08935, OJAG_RS08940, and OJAG_RS08945), *Microbacterium paraoxydans* (SAMN04489809_1122, SAMN04489809_1123, SAMN04489809_1124, SAMN04489809_1125, SAMN04489809_1126, and SAMN04489809_1127), *Agromyces subbeticus* (H521_RS21795, H521_RS21800, H521_RS0106365, H521_RS0106370, H521_RS21805, and H521_RS0106380), *Arthrobacter crystallopoietes* (D477_RS18370, D477_RS18375, D477_RS18380, D477_RS18385, and D477_RS18390, D477_RS18395), *Georgenia satyanarayanai* (DSZ44_RS04745, DSZ44_RS04750, DSZ44_RS04755, DSZ44_RS04760, DSZ44_RS04765, and DSZ44_RS04770), *Microbacterium trichothecenolyticum* (RS82_RS03115, RS82_RS03120, RS82_RS03125, RS82_RS03130, RS82_RS03135, and RS82_RS03140), *Arthrobacter woluwensis* (C6401_RS03950, C6401_RS03955, C6401_RS03960, C6401_RS03965, C6401_RS03970, and C6401_RS03975), *Promicromonospora kroppenstedtii* (PROKR_RS13815, PROKR_RS13820, PROKR_RS13825, PROKR_RS13830, PROKR_RS13835, and PROKR_RS13840), *Cellulomonas cellasea* (Q760_RS04595, Q760_RS04600, Q760_RS04605, Q760_RS18340, Q760_RS04615, and Q760_RS04620), *Agromyces cerinus* (BUR99_RS12060, BUR99_RS12065, BUR99_RS12070, BUR99_RS12075, BUR99_RS12080, and BUR99_RS12085), *Agreia pratensis* (B9Y86_RS06900, B9Y86_RS06905, B9Y86_RS06910, B9Y86_RS06915, B9Y86_RS06920 and B9Y86_RS06925), *Microbacterium laevaniformans* (OR221_3062, OR221_3063, OR221_3064, OR221_3065, OR221_3066, and OR221_3067), *Arthrobacter stackebrandtii* (CVV67_17780, CVV67_17785, CVV67_17790, CVV67_17795, CVV67_17800, and CVV67_17805), *Paeniglutamicibacter gangotriensis* (ADIAG_RS03760, ADIAG_RS03765, ADIAG_RS03770, ADIAG_RS03775, ADIAG_RS03780, and ADIAG_RS03785), *Microbacterium trichothecenolyticum* (RS82_RS03115, RS82_RS03120, RS82_RS03125, RS82_RS03130, RS82_RS03135, and RS82_RS03140), *Arthrobacter livingstonensis* (CVV68_RS19330, CVV68_RS19335, CVV68_RS19340, CVV68_RS19345, CVV68_RS19350, and CVV68_RS19355), *Demequina lutea* (AOP76_RS09030, AOP76_RS09035, AOP76_RS09040, AOP76_RS09045, AOP76_RS09050, and AOP76_RS09055), *Zhihengliuella halotolerans* (CUR88_RS12685, CUR88_RS12690, CUR88_RS12695, CUR88_RS12700, CUR88_RS12705, and CUR88_RS12710), *Paeniglutamicibacter antarcticus* (BN2261_RS08280, BN2261_RS08285, BN2261_RS08290, BN2261_RS08295, BN2261_RS08300, and BN2261_RS08305), *Janibacter melonis* (EEW87_RS00715, EEW87_RS00720, EEW87_RS00725, EEW87_RS00730, EEW87_RS00735, and EEW87_RS00740), *Microbacterium arborescens* (DOU46_RS02280, DOU46_RS02285, DOU46_RS02290, DOU46_RS02295, DOU46_RS02300, and DOU46_RS02305), *Agreia pratensis* (B9Y86_RS06900, B9Y86_RS06905, B9Y86_RS06910, B9Y86_RS06915, B9Y86_RS06920, and B9Y86_RS06925), *Agreia bicolorata* (TZ00_RS04480, TZ00_RS04485, TZ00_RS19215, TZ00_RS04495, TZ00_RS04500, and TZ00_RS04505), *Arthrobacter psychrochitiniphilus* (CVS30_RS02785, CVS30_RS02790, CVS30_RS02795, CVS30_RS02800, CVS30_RS02805, and CVS30_RS02810), *Microterricola pindariensis* (GY24_RS04745, GY24_RS04750, GY24_RS04755, GY24_RS04760, GY24_RS04765, and GY24_RS04770), *Microbacterium indicum* (H576_RS15860, H576_RS0112930, H576_RS0112935, H576_RS0112940, H576_RS0112945, and H576_RS15865), *Homoserinimonas sp.* (DL891_RS01870, DL891_RS01875, DL891_RS01880, DL891_RS01885, DL891_RS01890, and DL891_RS01895), *Cryobacterium levicorallinum* (SAMN05216274_11068, SAMN05216274_11069, SAMN05216274_11070, SAMN05216274_11071, SAMN05216274_11072, and SAMN05216274_11073), *Frigoribacterium sp.*(EDF18_RS14355, EDF18_RS14360, crtI, EDF18_RS14370, EDF18_RS14375, and EDF18_RS14380), *Cryobacterium luteum* (SAMN05216281_10883, SAMN05216281_10884, SAMN05216281_10885, SAMN05216281_10886, SAMN05216281_10887, and SAMN05216281_10888), *Cellulomonas carbonis* (N868_RS13600, N868_RS13605, N868_RS13610, N868_RS13615, N868_RS13620, and N868_RS13625), *Okibacterium fritillariae* (B5X75_RS14075, B5X75_RS14080, B5X75_RS14085, B5X75_RS14090, B5X75_RS14095, and B5X75_RS14100), *Glycomyces sambucus* (BLS99_RS13650, BLS99_RS13655, BLS99_RS13660, BLS99_RS13665, BLS99_RS13670, and BLS99_RS13675), *Krasilnikoviella flava* (B5Y66_RS20515, B5Y66_RS20520, B5Y66_RS20525, B5Y66_RS20530, B5Y66_RS20535, and B5Y66_RS20540), *Actinotalea ferrariae* (N866_01505, N866_01510, N866_01515, N866_01520, N866_01525, and ubiA), *Lysinimicrobium soli* (AOM04_RS11780, AOM04_RS11785, AOM04_RS11790, AOM04_RS11795, AOM04_RS11800, and AOM04_RS11805), *Luteimicrobium subarcticum* (CLV34_RS08275, CLV34_RS08280, CLV34_RS08285, CLV34_RS08290, CLV34_RS08295, and CLV34_RS08300), *Promicromonospora kroppenstedtii* (PROKR_RS13815, PROKR_RS13820, PROKR_RS13825, PROKR_RS13830, PROKR_RS13835, and PROKR_RS13840), *Tersicoccus phoenicis* (BKD30_RS05860, BKD30_RS05865, BKD30_RS05870, BKD30_RS05875, BKD30_RS05880, and BKD30_RS05885), *Sinomonas humi* (LK10_RS09295, LK10_RS09300, LK10_RS09305, LK10_RS09310, and LK10_RS09315), *Pseudarthrobacter phenanthrenivorans* (RM50_RS01675, RM50_RS01680, RM50_RS01685, RM50_RS01690, RM50_RS01695, and RM50_RS01700), *Acaricomes phytoseiuli* (C501_RS0107225, C501_RS0107230, C501_RS0107235, C501_RS0107240, C501_RS0107245,and C501_RS0107250), *Leucobacter musarum* (AMS67_RS10795, AMS67_RS10800, AMS67_RS10805, AMS67_RS10810, AMS67_RS10815, and AMS67_RS10820), *Ornithinimicrobium pekingense* (K330_RS19765, K330_RS0107130, K330_RS19770, K330_RS19775, K330_RS0107145, and K330_RS0107150), *Citricoccus sp.* (CITRI_RS16000, CITRI_RS0102550, CITRI_RS0102555, CITRI_RS16005, CITRI_RS0102565, and CITRI_RS0102570) and *Arthrobacter arilatensis* (AARI_13710, AARI_13720, AARI_13730, AARI_13740, AARI_13760, and AARI_13750)^8^ are very similar have a similar size and show the same organization as in the genome of *K. rhizophila*. In *Corynebacterium glutamicum* (cg0723, cg0721, cg0720, cg0719, cg0718, and cg0717)^9^ and *Corynebacterium efficiens* (HMPREF0290_1086, HMPREF0290_1088, HMPREF0290_1089, HMPREF0290_1090, HMPREF0290_1091, and HMPREF0290_1092)^10^ the decaprenoxanthin producing gene cluster is similar in size and organization to that of *K. rhizophila* except for the insertion of an unrelated gene cg0722 between *crtE* and *crtB* in *C. glutamicum* or HMPREF0290_1088 in *C. efficiens*. Also, in *Corynebacterium glutamicum*, an additional carotenoid cluster (NCgl0600, NCgl0598, NCgl0597, NCgl0596, NCgl0595, and NCgl0594) in also present in the genome. In *Kytococcus sedentarius*, the genes responsible for carotenoid production (Ksed_13840, Ksed_13830, Ksed_13820, Ksed_13810, Ksed_13800) are arranged in the same cluster while Ksed_16070, which encodes for geranylgeranyl pyrophosphate synthase is located elsewhere in the genome^11^.

In *Brevibacterium mcbrellneri,* the genes responsible for carotenoid production (HMPREF0183_0793, HMPREF0183_0794, HMPREF0183_0795, HMPREF0183_0796, and HMPREF0183_0797) are located in the same cluster while HMPREF0183_0437, which encodes for polyprenyl synthetase is located elsewhere in the genome.

In *Beutenbergia cavernae,* the genes responsible for carotenoid production (Bcav_3492, Bcav_3491, Bcav_3490, Bcav_3489, Bcav_3488) are located in the same cluster while Bcav_0970, which encodes for polyprenyl synthetase is located elsewhere in the genome^12^.

In *Brachybacterium faecium* (Bfae_04470, Bfae_04440, Bfae_04430, Bfae_04420, Bfae_04410, and Bfae_04400)^13^ the carotenoid biosynthetic cluster is similar in size and organization to that of *K. rhizophila* except for the insertion of two unrelated genes Bfae_04460 and Bfae_04450 between Bfae_04470 and Bfae_04440.

In *Cellulomonas flavigna* (Cfla_2888, Cfla_2889, Cfla_2890, Cfla_2891, and Cfla_2892)^14^ all the genes responsible for carotenoid production are present except for Cfla_2893 which is a likely pseudogene because of frame-shift mutation.

Decaprenoxanthin was the first C50 carotenoid discovered from *Flavobacterium dehydrogenan*s (now known as *Agromyces mediolanus*^15^).

Many bacteria including *Agromyces mediolanus*^15^, *Aureobacterium sp.*^16^, *Arthrobacter glacialis*^17^, *Arthrobacter arilatensis*^18^, *Cellulomonas biazotea*^19^, and *Corynebacterium glutamicum*^20^ are known to produce decaprenoxanthin.

